# Human bone marrow milieu identifies a clinically actionable driver of niche-mediated treatment resistance in leukaemia

**DOI:** 10.1101/2021.06.18.448490

**Authors:** Deepali Pal, Helen Blair, Sophie Boyd, Angel Hanmy Sharon, Salem Nizami, Asmida Isa, Melanie Beckett, Ryan Nelson, Aaron Wilson, Mankaran Singh, Shalini Sankar, Ricky Tirtakusuma, Nakjang Sirintra, Carly Knill, Andrew Fuller, Hesta McNeill, Lisa Russell, Claire Schwab, Peixun Zhous, Paul Sinclair, Jonathan Coxhead, Andrew Filby, Christina Halsey, James M. Allan, J. Christine Harrison, Anthony Moorman, Heidenreich Olaf, Josef Vormoor

**Author notes:** joint first (Deepali Pal, Helen Blair, Heidenreich Olaf, Josef Vormoor). joint last Affiliations.

## Abstract

Leukaemia cells re-program their microenvironment to provide proliferation support and protection from standard chemotherapy, molecularly targeted therapies, and immunotherapy. Although much is becoming known about molecules that drive niche-dependent treatment resistance; means of targeting these in the clinics has remained a key obstacle. To address this challenge, we have developed human induced pluripotent stem cell engineered niches *ex vivo* to reveal insights into druggable cancer-niche dependencies. We show that mesenchymal (iMSC) and vascular niche-like (iANG) cells support *ex vivo* proliferation of patient-derived leukaemia cells, impact dormancy and mediate therapy resistance. iMSC protected both non-cycling and cycling blasts against dexamethasone treatment while iANG protected only dormant blasts. Leukaemia proliferation and protection from dexamethasone induced-apoptosis was dependent on direct cell-cell contact and mediated by CDH2. To explore the therapeutic potential of disrupting this cell-cell interaction, we tested the CDH2 antagonist ADH-1 (previously in phase I / II for solid tumours) in a very aggressive patient-derived xenograft leukaemia mouse model. ADH-1 showed high *in vivo* efficacy. ADH-1/ dexamethasone combination therapy was superior to dexamethasone alone with no ADH1 conferred additional toxicity. These findings provide a proof-of-concept starting point to develop novel, potentially safer therapeutics that target niche-mediated cancer cell dependencies in haematological malignancies.

**Summary:** CDH2 mediated niche-dependent cancer proliferation and treatment resistance is clinically targetable via ADH-1, a low toxic agent that could be potentially repurposed for future clinical trials in acute leukaemia.

## INTRODUCTION

Treatment resistance remains a major obstacle in cancer management. Emerging evidence suggests that in addition to cell intrinsic mechanisms, factors such as the microenvironment are key in mediating cancer progression and escape from therapy(*1–4*). In direct contrast to the classical stem cell theory, which endows properties of quiescence and self-renewal to a rare hierarchical population of stem cells(*5*), recent studies attribute the microenvironment at least in certain tissues to function as mediators of stem cell self-renewal and differentiation(*6, 7*). An archetypal example is pancreatic ductal adenocarcinoma in which the tumor cells reprogram tumor-associated fibroblasts to provide a supportive and protective microenvironment(*8*).

Microenvironment conferred treatment resistance is a key impediment in treating blood cancers given leukaemic cells have a broad repertoire of tools to communicate with neighboring cells. These include direct cell-cell contact, tunneling nanotubes, exosomes and micro-vesicles, hormones and other soluble messenger molecules(*9–11*).

In addition, it is now well documented that leukaemia cells evolve their surrounding microhabitat and this dynamism not only enhances malignant propagation but also provides a safe haven against chemotherapy(*12, 13*). Leukaemic cells hijack the communication with bone marrow (BM) stroma and reprogram their microenvironment to survive therapy(*2, 13*). This communication is driven by molecular programmes such as IZKF1(IKAROS) deletions. IZKF1 deletions have been shown to induce expression of adhesion molecules, mediating strong adhesion to niche cells (including mesenchymal stem cells), integrin signaling and subsequently therapy resistance(*14, 15*).

Means of directly drugging cell-cell contact dependent treatment resistance with safe therapeutic agents are lacking. Key milestones in developing tractable *ex vivo* models for leukaemia niche interaction had been made by demonstrating that direct contact of acute lymphoblastic leukaemia (ALL) blasts with MSC in cell culture facilitates survival and limited proliferation *ex vivo*(*16–18*). However, improved and experimentally accessible models are needed for in-depth scrutiny of this intricate, multicomponent and continually evolving interaction. Using the complex BM as a paradigm, we micro-engineered human niche constituent cell types to define clinically exploitable cancer-niche interactions.

N-cadherin (CDH2) is a calcium-dependent transmembrane cell adhesion molecule known to regulate stem cell fate and proliferation(*19*). The cytoplasmic domains of N-cadherin bind to β-catenin as a linker to the actin cytoskeleton and association of N-cadherin with the cytoskeleton is necessary for stabilization of cell –cell adhesion(*20*). Cadherins are thought to play a crucial role and a potential target in the cell-cell contact of many tumor cells with their microenvironment. For CML, it has been shown that the N-cadherin/β-catenin complex is involved in mediating MSC-mediated resistance to tyrosine kinase inhibitors(*21*). Another study has shown that cordycepin, an agent with questionable stability in animal models prolongs survival in a CML cell line derived mouse model most likely via suppression of CDH2. Data on childhood ALL are limited. In ALL/t(1;19), expression of the oncogenic fusion protein E2A-PBX1 leads to overexpression of Wnt16. Via β-catenin, this mediates overexpression of N-cadherin and induction of cell-cell adhesion(*22*). The role of N-cadherin as a clinically actionable therapeutic target to disrupt malignant propagation and niche-mediated treatment response remains unexplored.

Here, we validate CDH2 as a druggable target in acute lymphoblastic and myeloid leukaemia. Our study highlights the opportunity to clinically repurpose ADH-1 (Exherin™), a low toxic drug that disrupts N-cadherin interaction. ADH-1 received orphan drug status in 2008 from the FDA and has previously been tested as an antiangiogenic agent in solid tumors in phase I/II trials(*23–25*).

## RESULTS

### Bone marrow induced pluripotent stem cell (BM-iPSC) derived bone marrow milleau support human haematopoietic cells *ex vivo*

In order to model the human leukaemia niche *ex vivo,* we re-programed primary BM mesenchymal stroma cells to pluripotency. This provided a replenishable and well defined source of BM constituent cells that represent both the mesenchymal stem and vasculature niche-like cells. Sendai virus is a highly efficient approach most commonly utilized for pluripotent reprograming however there are limitations to this technique(*26*). Most RNA-based approaches require repeat transfections due to reprogramming factor mRNA degradation(*27*). In light of this we adopted an RNA replicon reprogramming technology(*26*) that uses POU5F1, KLF4, SOX2 in combination with GLIS1 thereby replacing MYC and consequently endorsing a re-programming technology that is both virus and oncogene-free. Through standardized xeno-free protocols (Fig. 1A) we engineered 13 immortalized BM-iPSC lines (Fig.S1). Microsatellite DNA fingerprinting against parental mesenchymal cells confirmed authenticity (Table S1) whilst gene expression profiling revealed up-regulation of the embryonic stem cell genes SOX2, NANOG, GDF3, TERT, DNMT3B, CDH1, POU5F1 and ZFP42 (Fig. S 2.A.). BM-iPSC exhibited a pluripotent stem cell morphology and expressed the embryonic stem cell and pluripotency markers alkaline phosphatase, POU5F1, SOX2, SSEA4 and TRA-1-60 (Fig. S2.B,C). *In vitro* embryoid bodies (Fig. S2.D,E) and *in vivo* teratomas showing ectodermal, mesodermal and endodermal germ layer differentiation (Fig. 1. B.) confirmed the pluripotent nature of BM-iPSC at a functional level.

**Figure 1.**
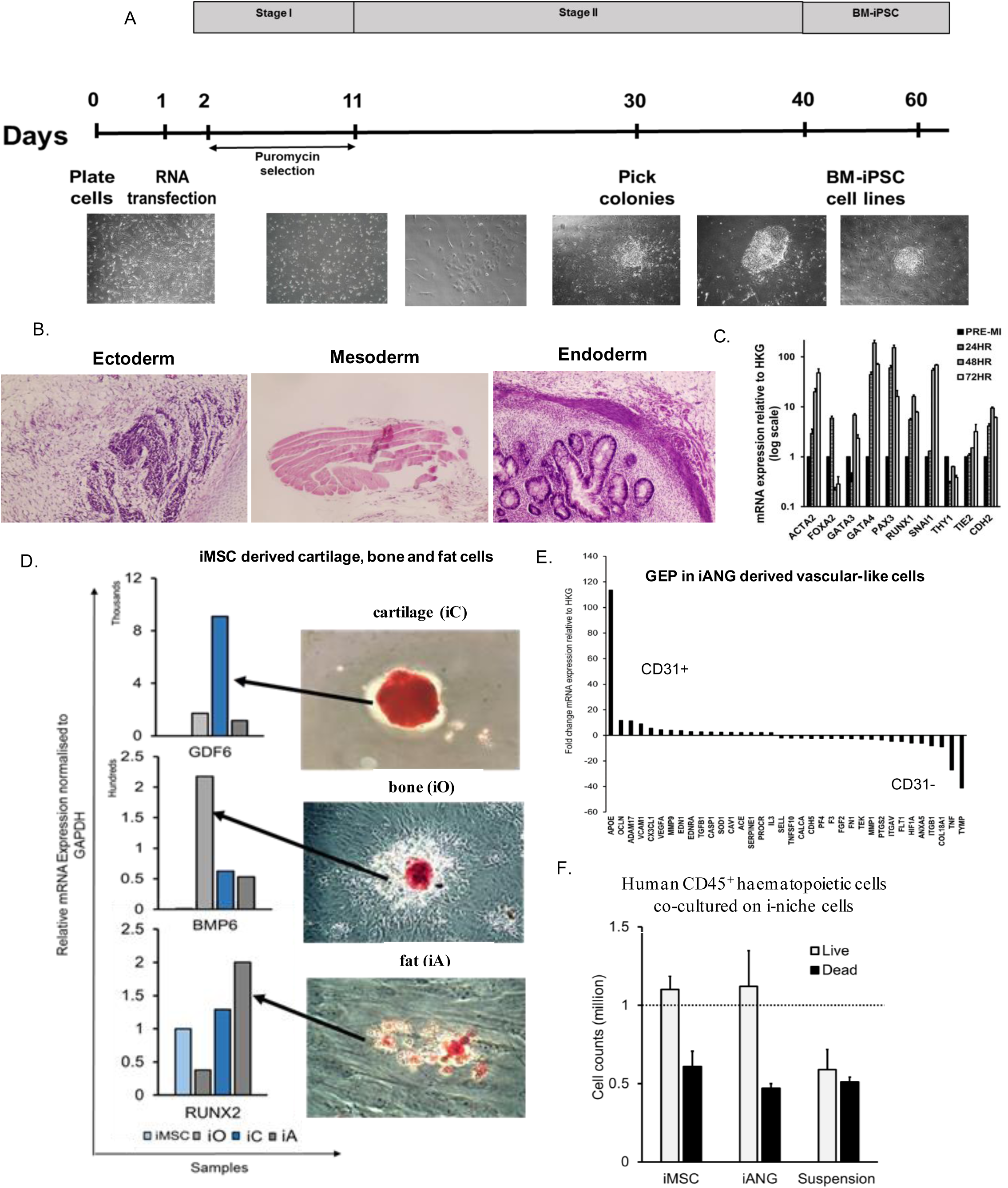
BM-iPSC derived bone marrow milieu support human haematopoietic cells ex vivo. A. Schema for synthetic RNA based re-programming using pluripotent transcripts POU5F1-SOX2-KLF4, GLIS1 B. H&E staining of BM-iPSC-derived teratomas representing the three embryonic lineages. C. mRNA expression of BM-iPSC cells during mesodermal differentiation: pre mesoderm induction and at 24, 48 and 72 hours after mesoderm induction. D. GDF6, BMP6 and RUNX2 expression in i-MSC derived cartilage/chondrocytes, bone/osteoblasts and fat/adipocytes cells (iC, iO and iA). Immunohistochemical staining demonstrating Safranin O, Alizarin Red and Oil Red O staining in iC, iO and iA respectively. E. Gene expression profile (GEP) in iANG containing representative vascular cells such as CD31+ endothelial cells [25%] and CD31- perivascular cells [75%] in known proportions. CD31+ cells express endothelia- relevant markers such as APOE, OCLN, ADAM17, VCAM1 whereas CD31- cells express perivascular markers such as ANXA5, ITGB1, HIF1A and COL18A1 F. Cell counts of non malignant CD45^+^ haematopoietic cells extracted f rom human bone marrow and co-cultured on iMSC, iANG versus in niche-f ree suspension cultures over 7 days.

Next, mesenchymal (iMSC) and vascular niche-like (iANG) cells (together i-niche) were derived from BM-iPSC through a mesoderm intermediate. Within 72 hours of initiating mesoderm induction, tightly packed pluripotent cells with high nuclear to cytoplasmic ratio began to alter their morphology to form cobblestone clusters comprised of polygonal cells (Fig. S3.A). Gene expression profiling confirmed downregulation of pluripotent (Fig. S 3.B) and upregulation of mesodermal genes (Fig. 1C) thus corroborating directed differentiation of BM-iPSC into mesodermal lineage. Furthermore, we observed upregulation of WNT5A during this process. WNT5A is observed in human embryonic stem cell-derived mesoderm(*28*) and upregulation of this gene confirms lineage-specific directed differentiation of BM-iPSC (Fig. S3.C). We further differentiated these early mesoderm cells into iMSC and iANG that exhibited clearly distinct transcriptomic patterns with iMSC upregulating mesenchymal genes (Fig. S.4.A,B). Differentiation of iMSC into osteogenic, chondrogenic and adipogenic cells (Fig. 1.D, S4.C) further confirmed their mesenchymal stem or progenitor potential. iANG cells contained a population of CD31+ endothelia-like cells and CD31- perivascular-like cells (Fig. 1.E). We further confirmed that CD31+ cells upregulated expression of genes such as APOE which has been documented to be localized to endothelial cells *in vivo*(*29*), VCAM1, an endothelial cell surface glycoprotein(*30, 31*), CX3CL1, known to be produced by endothelial cell membranes(*32, 33*), and OCLN, a functional marker of endothelial cells linked to their ability of tube formation(*34*). We reveal that CD31- cells, on the other hand, expressed genes more closely associated with perivascular cells such as ANXA5(*35*), ITGB1(*36*) and HIF1A(*37*). To specify the role of iMSC and iANG in sustaining hematopoiesis, we isolated CD45+ cells from non-malignant human BM for co-culture on iMSC and iANG. Unlike microenvironment-free suspension cultures, both types of niche cells supported viability of human bone marrow derived hematopoietic cells (n=3 different donors) (Fig. 1F). Together these data show that primary mesenchymal stroma stem-cell re-programmed into BM-iPSC via a virus and C-MYC free RNA-based route are able to differentiate into mesenchymal stem cells and vascular BM niche like cells. In addition, both i-niche cell types successfully supported ex vivo survival of human blood cells.

### Niche–primed leukaemic cells upregulate CDH2

To further define the clinical relevance of i-niche cells in blood cancer, we evaluated and characterized their potential to re-create a microenvironment that would support survival, self-renewal and proliferation of malignant cells. Blasts from several patient-derived leukaemia samples (n=14 samples) proliferated on i-niche cells (Fig. 2B, Table S2). There was parity between iMSC and primary MSC in supporting ALL cells (Fig S5.A). We showed that direct niche contact was superior in supporting leukaemic proliferation compared to feeder-conditioned media (Fig. S5.B,C). Using FISH analysis and whole exome sequencing experiments we confirmed not only maintenance of the leukaemia but also of > 99% genomic complexity on both iMSC and iANG (Fig. S6).

**Figure 2.**
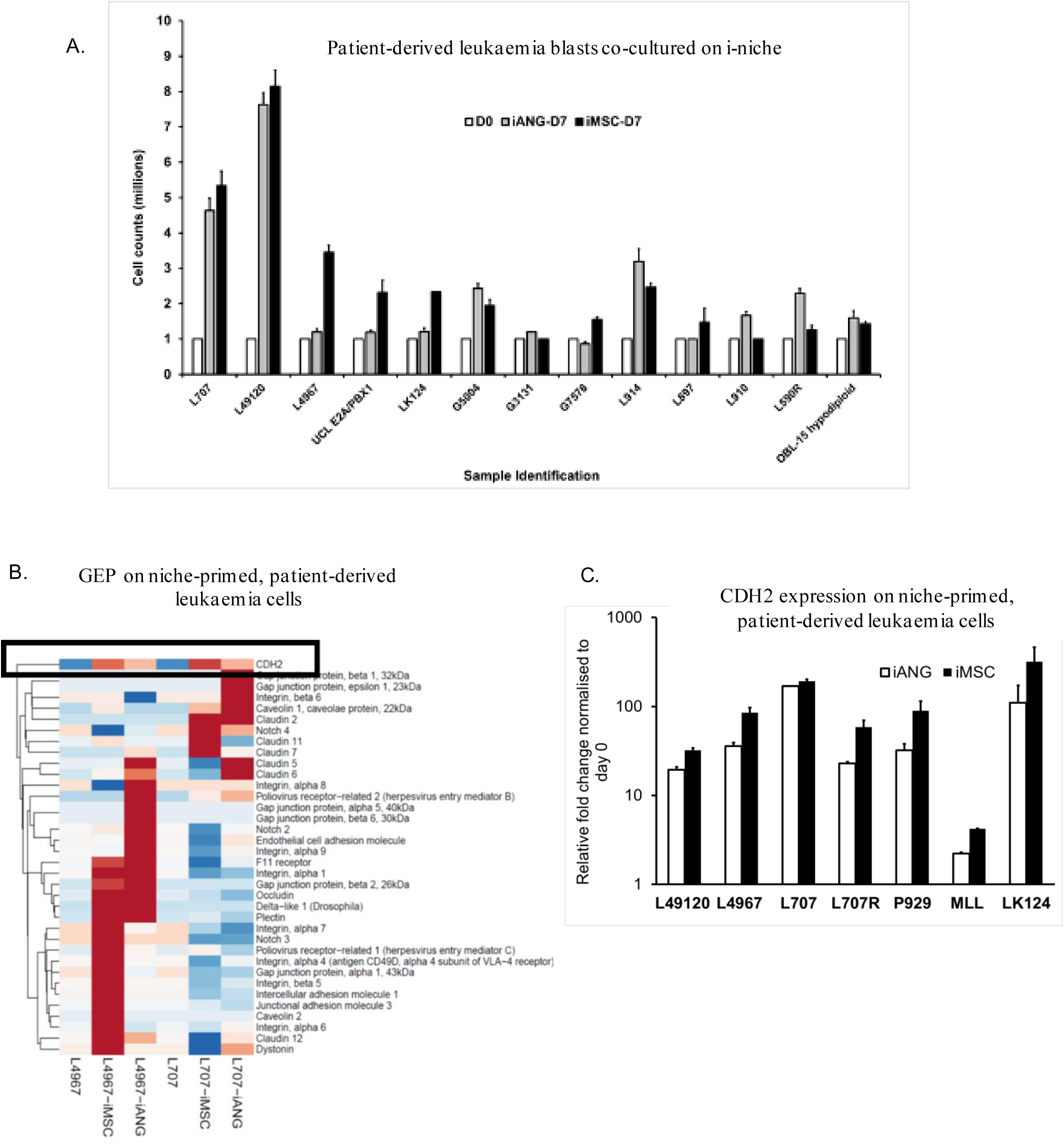
Niche – primed leukaemia cells upregulate CDH2. A. Cell counts of patient-leukaemic blasts on iMSC and iANG at diagnosis and relapse over a seven day period. B. Heatmap demonstrating gene expression profiling of niche primed patient-leukaemia samples [L707, L4967] shows consistent upregulation of CDH2 following a 7 day co-culture with iMSC and iANG C. CDH2 upregulation confirmed by qRT-PCR on 7 leukaemia samples following a 7 day co-culture with iMSC and iANG

In order to study the effects of the different i-niche cells on supporting lymphoid and myeloid cell types we co-cultured leukaemic cells from a patient with infant ALL/t(4;11) who initially presented with a CD34+CD19+CD33+CD15- immunophenotype but relapsed at 5 months with a myeloid CD34+CD19-CD33+CD15+ leukaemia. Both iMSC and iANG supported maintenance of CD34+CD19+ lymphoid leukaemic cells (lympho-permissive). In contrast some cells in suspension culture lost expression of both CD34 and CD19. On iMSC, the leukaemic blasts lost expression of the myeloid marker CD33+ (myelo-suppressive) while on iANG the blasts retained expression of CD33 with emergence of a population of CD15+ cells (myelo-permissive). (Fig. S7, Table S2)

In a previous study, we confirmed that cell-cell contact between primary BM mesenchymal stem cells and leukaemia cells plays a significant role in supporting proliferation of leukaemic blasts(*16*). Based on the known role of adherens junctions in cell-cell contact and cancer cell-niche communication we conducted gene expression profiling with a focus on adherens junction molecules using a combined approach of RNA sequencing, qPCR arrays and real-time qPCR experiments on iMSC and iANG primed patient-derived blasts. Analysis of blasts following a 7 day priming (co-culture) on the i-niche cells showed upregulation of several genes relating to adherens junction, WNT and β-Catenin pathway genes (Fig. S8). In line with our observation that niche-mediated leukaemia survival and proliferation is regulated by direct cell contact, we also saw upregulation of several cell-cell junction and cell adhesion molecules on leukaemic blasts that were co-cultured with i-niche over seven days. These blasts were harvested from co-cultures and following cell separation through filtration subjected to gene expression profiling experiments. We found consistent upregulation of cell adhesion molecule CDH2 in i-niche primed blasts across two patient leukaemia samples (Fig. 2C). We further validated upregulation of CDH2 on a total of 6 diagnostic and 1 relapse patient samples (Fig. 2D). In summary, these data show that BM-iPSC derived niche cell types support leukaemia cells and blasts primed by the i-niche cells upregulate CDH2 expression.

### Under dexamethasone treatment, CDH2 is upregulated by iMSC-primed cycling cells

We next studied the role of CDH2 in niche-mediated cancer cell quiescence and proliferation. DNA labelling dyes allow isolation and tracking of dormant cells identified as the non/slow dividing and label retaining population(*38*). Cell generational tracing experiments were performed to compare patterns of leukaemic dormancy between the mesenchymal and vascular niche-like microenvironments. A patient with ALL/t(17;19) (Table S2) who initially presented with steroid-sensitive leukaemia but later relapsed with steroid-resistant disease (due to a homozygous deletion of the glucocorticoid receptor NR3C1) was used as a model to study the effects of dexamethasone in our i -niche system.

Distinct patterns of leukaemia quiescence and proliferation were detected on the two niche cell types (Figure 3.A.). iMSC only supported fast dividing blasts (label^low^). In contrast, nearly 50% of the total patient-derived blasts on iANG cells were non dividing cells (label^high^). Both iMSC- and iANG-primed blasts engrafted immunocompromised mice although iANG-primed blasts appeared to preferentially home to the murine BM and to a lesser degree to the spleen (Fig. 3B, Table S3). To further define the role of the different i-niche cells on leukaemic quiescence and proliferation, we extended our analysis to include cells from the matched relapse sample (Fig. 3C-3E). Cells from the diagnostic sample proliferated faster on iMSC while the relapse cells proliferated faster on iANG cells (Fig. 3C). Correspondingly, Hoechst/Pyronin Y staining experiments showed a 4 fold higher percentage of cells from the diagnostic sample in G0 on iANG niche cells (Fig. 3D). There was no difference in the percentage of cells from the relapse sample in GO or in the cell cycling pattern when cultured on either iMSC or i-ANG (Fig. 3D & E).

**Figure 3.**
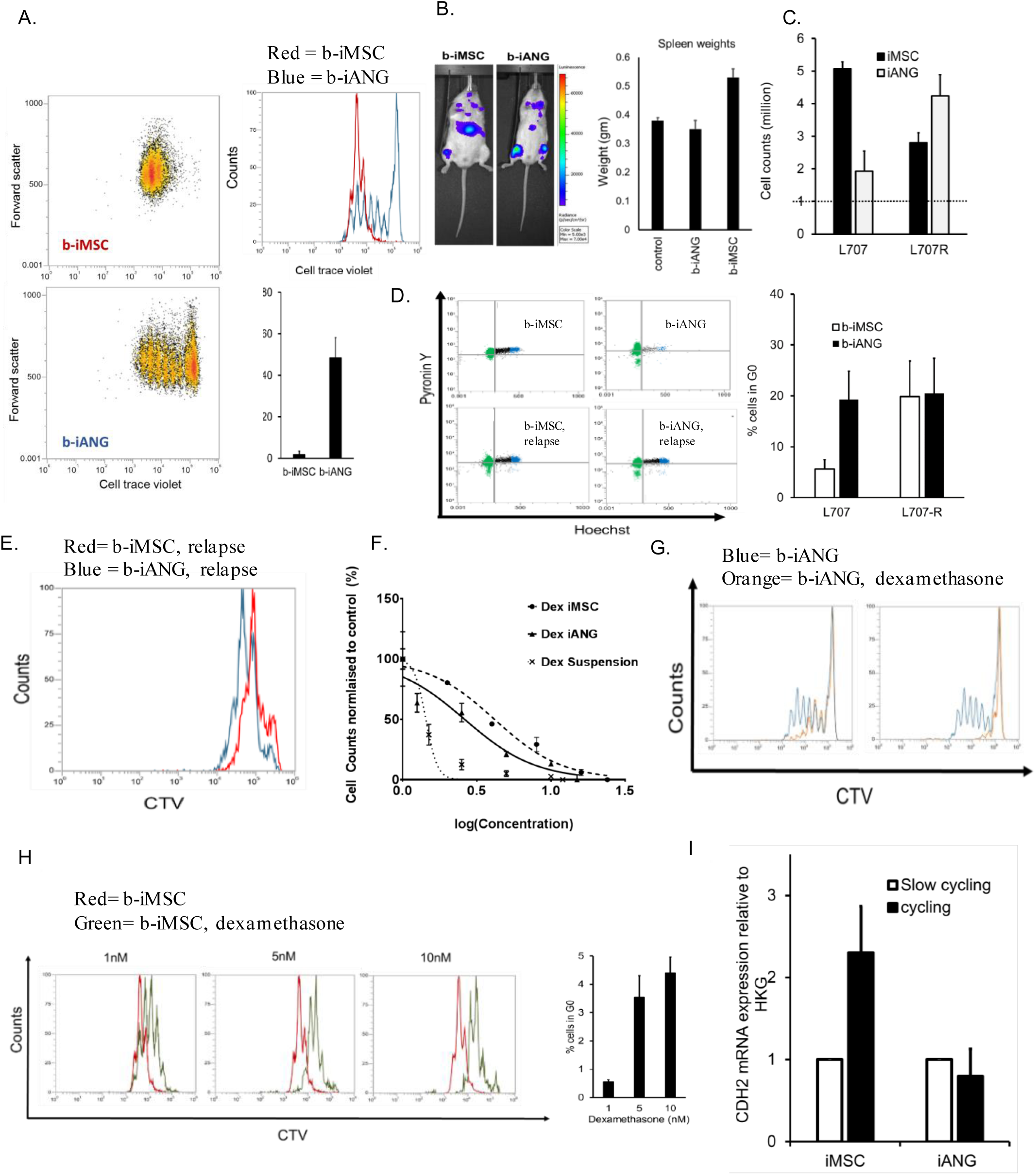
Under Dexamethasone treatment pressure CDH2 is upregulated by iMSC-primed cycling cells. A. Dot plots show fast cycling and slow cycling iMSC primed blasts [b -iMSC, red] and iANG primed blasts [b-iANG, blue] at Day 7 f rom a patient-leukaemia (L707) sample at diagnosis. Graph shows % slow cycling blasts on iMSC and iANG. B. Total fluorescence intensity of luciferase-tagged niche-primed patient leukaemic blasts transplanted in immunocompromised mice [intrafemoral transplants, n= 3 independent experiments, 1 representative experiment shown]. The column graph depicts spleen weights (harvest at 4.5 weeks following injection) in mice transplanted with patient-blasts (L707) at diagnosis (control) and following a 7 day co-culture on iMSC (b-iMSC) and iANG (b-iANG) C. Cell counts of a diagnostic and matched relapse sample following co-culture on iMSC and iANG. D. Hoechst-pyronin Y analysis [dot plot] of patient leukaemic blasts on iMSC [lef t panel] and iANG [right panel] in patient leukaemic blasts at diagnosis [top panel] and relapse [bottom panel]. Graph shows percentage cells in G0 on iMSC [b -iMSC] and on iANG [b-iANG] at diagnosis [L707] and relapse [L707-R]. E. Fast and slow dividing niche primed blasts f rom relapse sample on iMSC [red] and iANG [blue] at Day 7. CTV = Cell trace violet dye F. Dexamethasone dose response (nM) curve of patient leukaemia cells (treated for 7 days) in niche-free suspension culture and on iMSC and iANG G. Histogram shows cell generatio nal curve of untreated [blue] and treated cells [orange]. H. Cell generation curves of patient leukaemic cells untreated [red] and treated [green]. Column graph shows % slow cycling blasts on iMSC under Dexamethasone treatment. I. CDH2 expression under dexamethasone pressure in slow cycling and cycling/fast cycling blasts relative to HKG. Blasts were sorted using f low cytometry following seven day treatment with 5nM Dexamethasone. N=3 for 1 patient sample, L707 shown here. Data for 3 additional patient samples are included in S25

To study niche-mediated resistance, we repeated the cell division tracing under treatment pressure. In compliance with the clinical and molecular data, cells from the diagnostic sample were sensitive to dexamethasone while relapse blasts showed no response (Fig. S9). Dose response curves demonstrated reduced sensitivity against dexamethasone on both types of i-niche cells as compared with the niche-free suspension cultures (Fig. 3F). On iANG cells, dexamethasone treatment actively killed the dividing blasts, mainly leaving a non-dividing label^high^ population intact (Fig. 3G). On iMSC, treatment caused the cell division curve to shift to the right depicting cell populations that were dividing more slowly and with the emergence of only a small (5%) non dividing population (label^high^) (Fig. 3H). Unlike iANG cells, iMSC cells facilitated survival of slower dividing ALL blasts under dexamethasone treatment suggesting that treatment resistance is unlikely to be attributed to dormancy alone. Subsequently we re-visited the role of CDH2 in proliferation and treatment resistance. We saw in four patient samples that fast dividing, label^low^ iMSC- primed blasts that survive under dexamethasone pressure expressed higher levels of CDH2 (Fig. 3I, S10). These results suggest that CDH2 plays a direct role in mediating niche-dependent leukaemia proliferation in blasts that are resistant to treatment with dexamethasone.

### CDH2 drives leukaemia proliferation and reduces sensitivity against dexamethasone

In order to validate the function of CDH2, we performed RNAi knockdown experiments on both cancer cells and i-niche cells. CDH2 knockdown in 4 different leukaemia cell lines (Fig. 4A., Fig. S11) resulted in reduced proliferation in niche-free suspension cultures (Fig. 4B, 4C). Moreover, CDH2 knockdown resulted in downregulation of a range of cancer associated gene signatures (Fig. S12) including key oncogenic pathways, such as JAK-STAT, prolactin, chemokine and ErbB signaling (Figure S12) as well as modulation of several genes associated with leukaemogenesis and transcription and chromatin remodeling factors (Fig S13).

**Figure 4.**
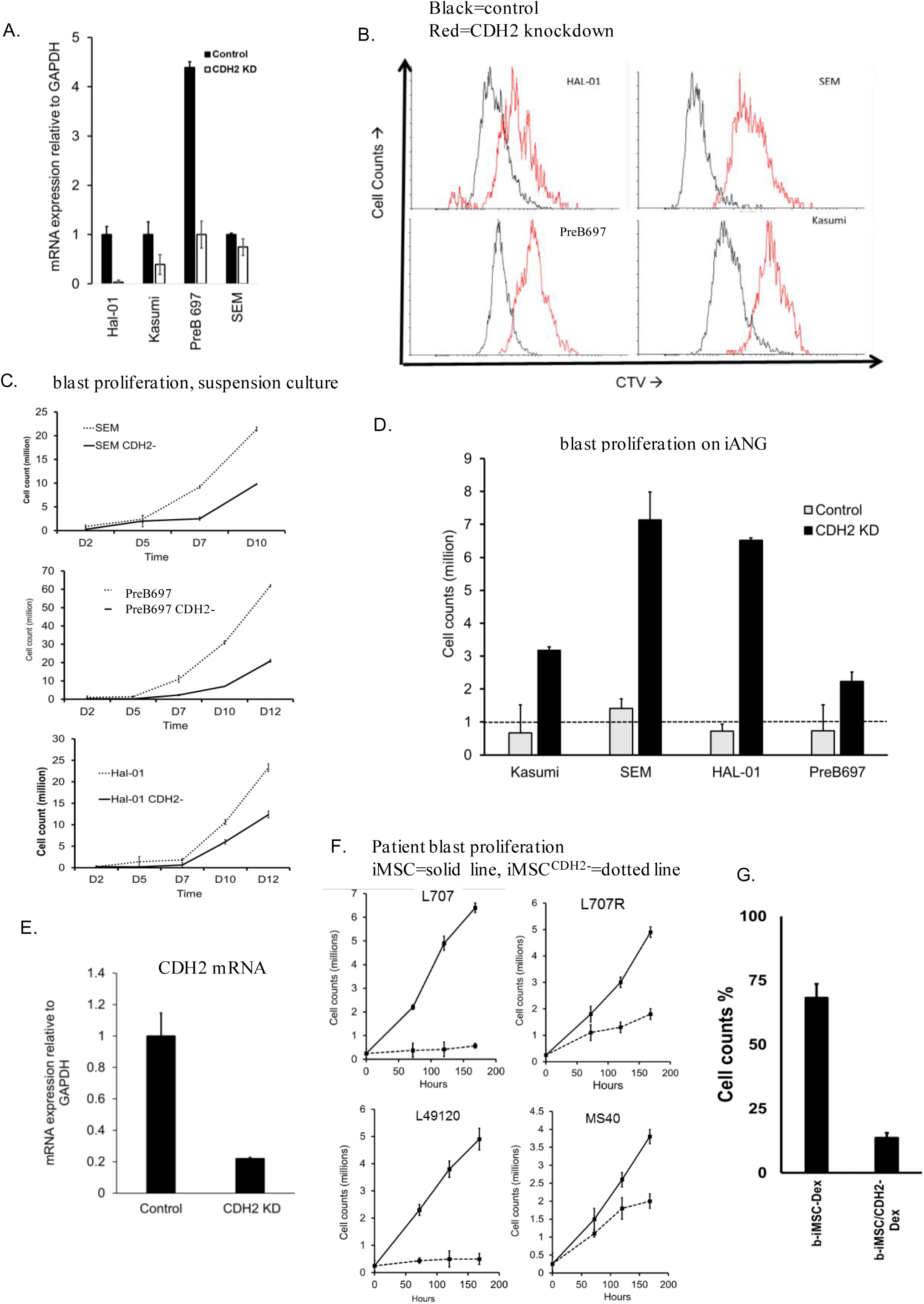
CDH2 drives leukaemia proliferation and reduces sensitivity against Dexamethasone. A. CDH2 levels in leukaemia cell lines following lentiviral knockdown. Control = nonsense siRNA/empty vector B. Cell generational tracing curves using the dye cell trace violet (CTV) in 4 different leukaemia cell lines following CDH2 knockdown. Black = empty vector control. Red = CDH2 knockdown C. Leukaemic cell proliferation in three different acute lymphoblastic leukaemia cell lines following CDH2 knockdown [against empty vector control]. D. Cells counts of CDH2 knockdown and empty vector control cell lines on iANG over 5 days, n=2. Dashed line indicates a starting cell count of 1 million cells. Feeder dependence was achieved by conducting co-cultures in the absence of FBS and at a reduced leukaemia cell density of 10,000 cells/ml. Under these altered culture conditions the leukaemia cells failed to survive on iMSC. E. CDH2 mRNA levels in control iMSC and CDH2 knockdown iMSC (iMSCCDH2-). F. Cell counts of three different patient leukaemia samples on iMSC (solid line) and iMSC^CDH2-^(dotted line). G.% cell counts (with respect to untreated control) of patient leukaemia cells on iMSC^CDH2-^ with and without 5 nM dexamethasone.

CDH2 knock-down leukaemia cells, co-cultured under modified culture conditions to facilitate niche-dependence, failed to survive on iMSC cells and showed reduced proliferation on iANG (Fig. 4D). Conversely, CDH2 knockdown in iMSC (Fig. 4E) reduced their ability to support the proliferation of three diagnostic and one relapse patient-derived leukaemia samples (Figure 4.F). Importantly, the leukaemic blasts demonstrated a 3 fold higher sensitivity to dexamethasone on iMSC^CDH2-^ cells (Fig. 4.G) These data suggest that BM mesenchymal stem cells mediate their leukaemia supportive effect via heterologous cancer-niche interactions through CDH2-CDH2 binding and signaling.

### CDH2 antagonist ADH-1 shows high *in vitro* efficacy in patient-derived leukaemia cells

ADH-1 is a small, cyclic pentapeptide with the formula N-Ac-CHAVC-NH2 that competitively blocks the action of CDH2. In preclinical models it has antiangiogenic properties in disrupting tumor vasculature and inhibiting tumor growth. The compound has been in Phase I / II trials for advanced solid malignancies(*23–25*) and received orphan drug status from the FDA in 2008. Its efficacy in blood cancers remains unknown. We applied our i-niche co-culture platform and demonstrated sensitivity to ADH-1 in 15 different patient-derived leukaemia samples (Fig. 5A; Figure S14, Table S2). ADH-1 doses used throughout this study are at par with plasma level concentrations that have been achieved in solid tumor trials(*39*). We found that ADH-1 treatment showed maximum efficacy when the leukaemic cells were in direct contact with the niche as opposed to transwell cultures (Fig. 5C-D). In keeping with this, ADH-1 treatment in cancer-niche co-cultures increased leukaemia cell death as evidenced by increased Annexin V, PI staining (Figure 5.E).

**5.**
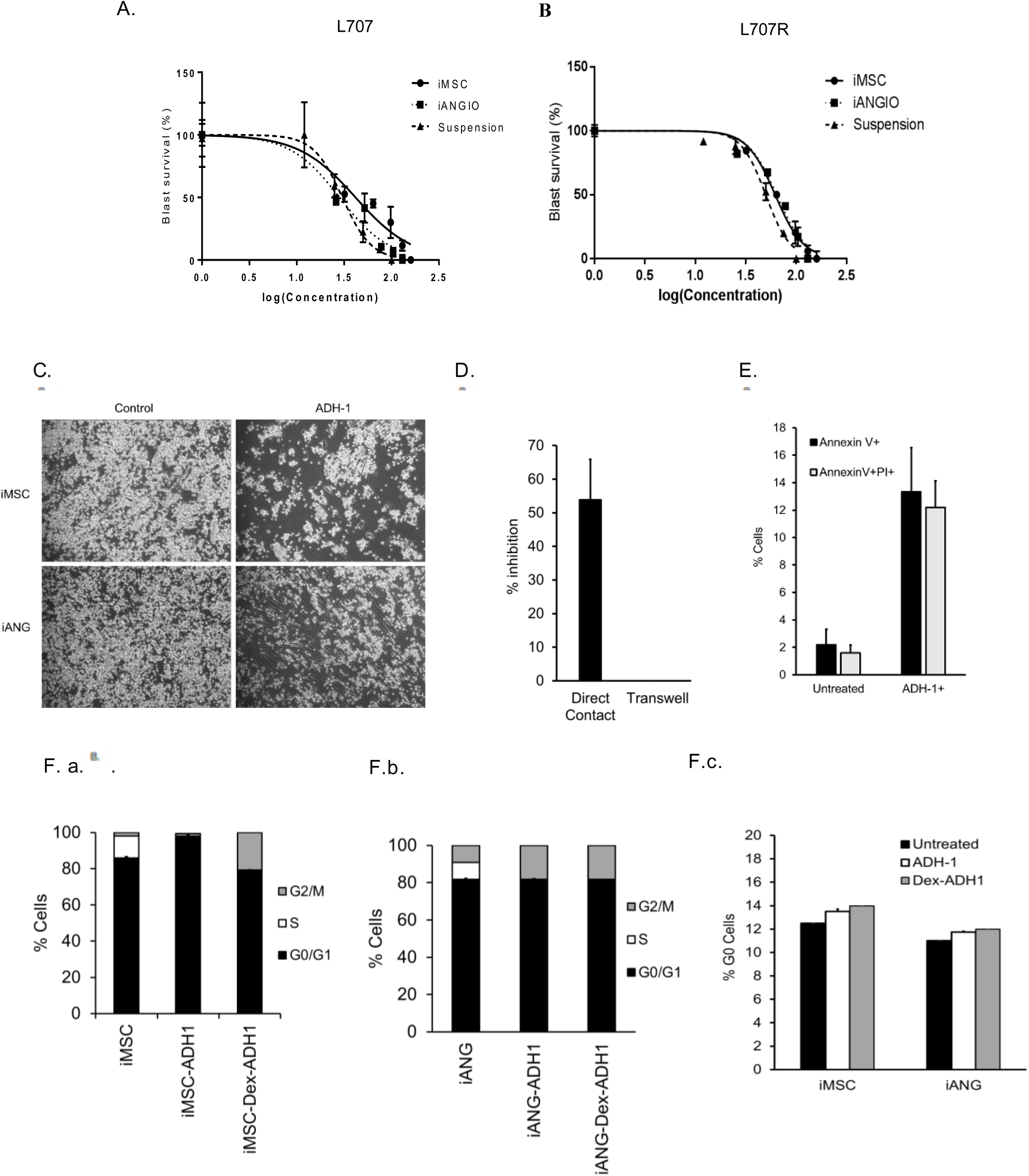
CDH2 antagonist ADH-1 a repurposed compound is identified to show high efficacy on a wide range of patient derived leukaemia cells. A. ADH-1 dose response curves in patient leukaemia samples f rom a patient at diagnosis and B. relapse. C. Adherent patient blasts on iMSC and iANG following treatment with ADH-1. D. % inhibition (cell counts) of blasts following ADH-1 treatment on direct contact cultures [iMSC] and in transwell cultures E. Annexin V PI f low cytometry analysis in patient blasts following treatment with ADH-1. F a-c. RNA and DNA content analysis using f low cytometry in primary blasts following treatment with ADH-1 in a. iMSC and b. iANG co-cultures Column graphs show % cells in S and G0 phase following treatment with ADH-1. c. % G0 cells in co-cultures following treatment with ADH-1

To further investigate the effect of this compound on cells that survive under treatment pressure we performed live cell cycle and G0 analysis on patient-blasts at relapse. We found that ADH-1 treatment slowed down proliferation of the leukaemic blasts as evidenced by reduced number of cells in the S phase. Furthermore, there was no accumulation of cells in G0 (Fig. 5F). Taken together these data corroborate that ADH-1 is a promising candidate for the treatment of resistant relapse without inducing drug resistant quiescent cells.

Despite the recent improvements in targeted therapeutics single agent treatment has been associated with emergence of treatment resistant cancer clones(*40, 41*). Combinatorial drug treatment is a central principle in anti-cancer therapy not only to enhance efficacy through drug synergies but most importantly to prevent emergence of treatment resistance. Drug combination assays with dexamethasone and ADH1 using a total of four different patient-derived leukaemia samples (Fig. 6A-F) showed synergistic interaction as analyzed by the Bliss Independence model. Comprehensive drug matrix analyses (Fig. 6F-G) demonstrated synergy for ADH-1 in combination with clinically relevant concentrations of dexamethasone achieving ZIP synergy scores of >10 on both iMSC and iANG. Taken together these data reveal a new way of clinically targeting niche-mediated leukaemia treatment resistance using the CDH2 antagonist ADH-1.

**Figure 6.**
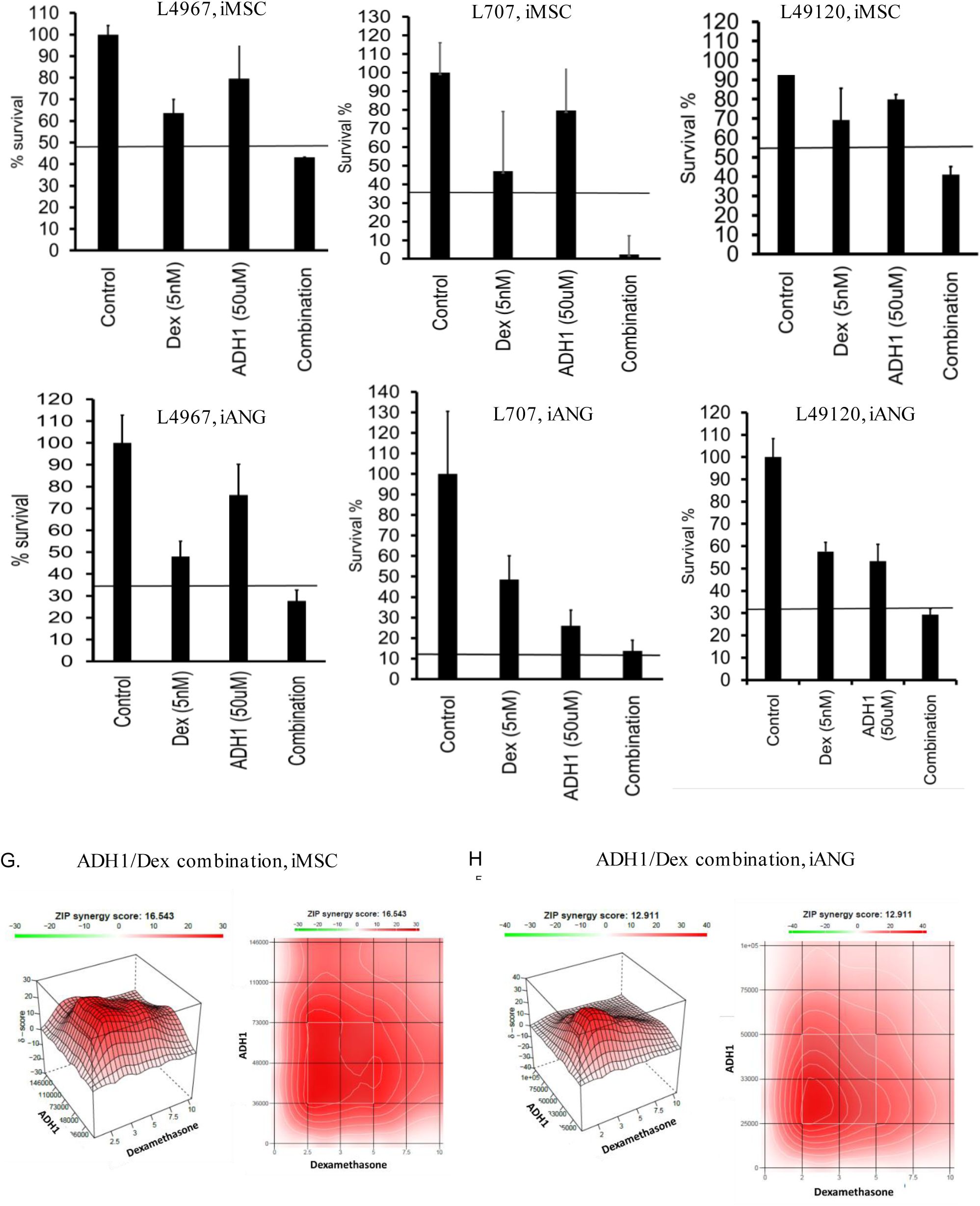
ADH-1 demonstrates in vitro synergy in combination with Dexamethasone. % survival following treatment with Dexamethasone, ADH-1 and combination in three different patient samples over seven days: A. L4967 B. L707D and C. L49120. On iMSC and D-F on iANG. Horizontal line depicts the expected combined effect as per the Bliss independence model. G. Synergy landscapes (3D and 2D synergy maps) and ZIP synergy scores of ADH-1/Dex on primary patient blasts (L707) on iMSC and H. iANG co-cultures over a seven day period.

### 6. ADH-1 shows high *in vivo* efficacy in a very aggressive leukaemia patient derived xenograft (PDX) model

In order to validate the function of CDH2 *in vivo*, we interrogated leukaemia initiation and propagation in our PDX model(*42–44*). We transplanted luciferase-tagged leukaemic blasts from a clinical sample of very high risk acute lymphoblastic leukaemia (L707, Table S1) directly into the bone marrow of immunodeficient mice. We monitored leukaemic engraftment into the mouse bone marrow via bioluminescence and we confirmed successful engraftment through immunohistochemistry staining of mouse bone marrow with human CD19 (a lymphoid cell marker) (Fig. S15). In an initial pilot we treated mice with a combination of ADH-1 and dexamethasone to determine a non-toxic dose and schedule for further study (Fig. S16A). This small-scale dose escalation pilot study indicated that ADH1 and dexamethasone in combination significantly reduced leukaemic engraftment (Figure S16.B) thereby justifying further *in vivo* investigation of the combination treatment. We also found that ADH1/dexamethasone [ADH1 200mg/kg; dexamethasone 3mg/kg] delivered via intraperitoneal injection was well tolerated when administered 5 times weekly for 3 weeks with minimal weight loss. ADH1 dosing was based on previously published studies in mice(*45, 46*) and the dexamethasone dose chosen to replicate plasma concentrations achieved in ALL patients(*39, 47, 48*). We repeated the *in vivo* transplantation experiments using bioluminescent-tagged patient-derived ALL blasts and started drug dosing on day 6 following transplantation (Fig. 7A). By bioluminescence monitoring, ADH-1 alone showed a similar reduction in leukaemic progression as observed in mice treated with dexamethasone. More importantly, the ADH1/dexamethasone combination treatment profoundly reduced leukaemic engraftment compared to controls and single agent therapy. Through additional bioluminescent imaging (BLI) we demonstrated significantly lower overall signals compared to untreated controls at both weeks 2 and 3 of ADH1/dexamethasone therapy (Fig 7B-C). Confirming the imaging data, spleen sizes were significantly smaller in the ADH1/dexamethasone treated mice at the end of the study. We further showed that the proportion of leukaemic blasts in bone marrow and spleen was significantly less in ADH1/dexamethasone treated mice compared with mice from the dexamethasone and control groups (Fig 7D-E, S16.C). In keeping with our *in vitro* observations, the ADH1/Dexamethasone combination was most effective in the bone marrow (7D-E) suggesting that a key mechanism of action for ADH-1 was to disrupt CDH2-mediated blast-bone marrow niche interactions increasing sensitivity to dexamethasone.

**Figure 7.**
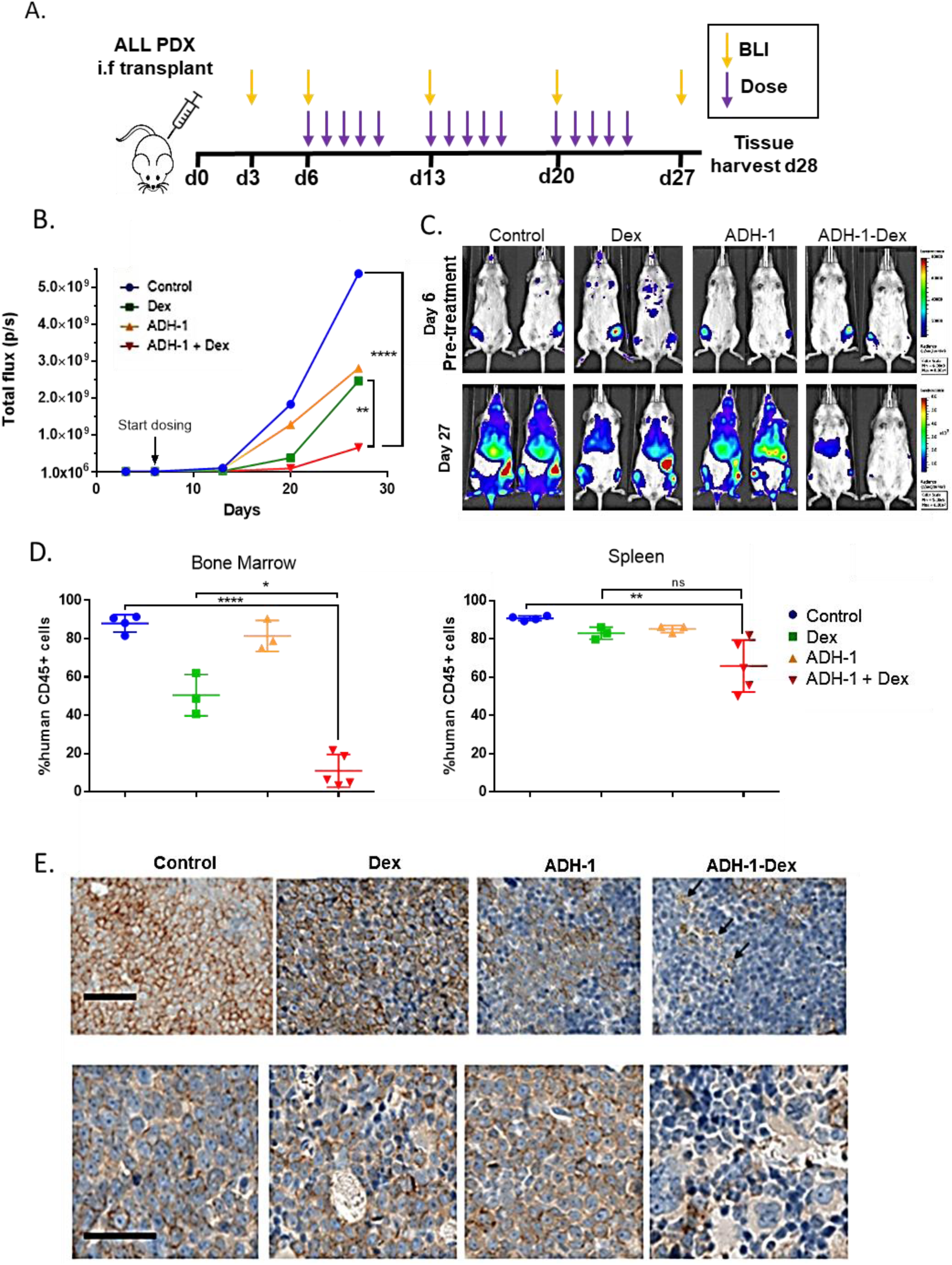
ADH-1 potentiates dexamethasone sensitivity *in vivo*. A. The PDX *in vivo* efficacy study design. Mice were dose interperitoneally with either, saline vehicle (control), 3mg/kg dexamethasone (Dex), 200mg/kg ADH-1 or ADH-1 Dex combined, 1x daily, 5x weekly for 3 weeks (15 doses). B. Mean whole-body total f lux measurements f rom bioluminescent imaging of each treatment group. C. Representative luminescence images of mice before and after treatment. Mice at each time point are show with identical luminescence scale for comparison. Leukaemic blasts are present in the femurs of all mic e at the start of treatment. Signal spreads to bone marrow sites, liver and spleen in control mice whereas signal is barely visible in ADH1-Dex controls. D. Leukaemic engraftment in harvested bone marrow and spleen measured by f low cytometry of labelled harvested cells. Human CD45+ cells are shown as a % total CD45+ cells (mouse + human cells). Lines indicate mean and SE, symbols for individual mice. ANOVA (GraphPad Prism), ns not significant, * p<0.05, **p<0.005, ****p<0.00005. E. Human CD19 Immunohistochemistry on sections of spleen and bone harvested f rom mice. Mice treated with ADH-1/Dex combination have few CD19 stained cells (brown staining at the cell membranes) and have areas of punctate staining indicative of cell debris (arrows). Scale bar = 50µm.

In summary, we found that treatment with only 15 doses of dexamethasone in the presence of ADH1 was more effective than dexamethasone alone at preventing leukaemic growth *in vivo*. This *in vivo* efficacy validated the use of our engineered preclinical model for the identification of exploitable niche targets.

## DISCUSSION

Treatment resistance and treatment toxicity are major clinical challenges that urgently need more attention. Dynamism of the leukaemic niche and its role in dormancy and treatment resistance is well documented(*2, 38, 49-51*). Indeed, standard chemotherapy primes cancer and its ambiance alike endowing cell intrinsic and non -cell-autonomous adaptations towards treatment resistance(*2, 52*). Recent concepts such as non-oncogene addiction(*40*), a phenomenon underpinning cancer cell survival through exaggerated functioning of non-mutated genes have emerged as promising solutions to prevent treatment resistance. Despite the significant impacts of the niche on cancer cell function; no druggable niche targets exist that can directly impact microenvironment mediated leukaemia biology. Indeed, current day treatment largely disregards the influence of the oncogenic microenvironment on malignant proliferation, self -renewal and treatment resistance.

To identify new and safer actionable targets against the leukaemic niche we require better preclinical models. This is a major challenge in hematological malignancies since most primary leukaemia cells do not proliferate once removed from the patient and their microenvironment. Consequently, there is a lack of models that allow scrutiny of the niche in a human cell-based setting(*16, 53*). Here we have shown that human pluripotent stem-cell engineered BM cells support *ex vivo* proliferation of patient-derived leukaemia cells. Furthermore, different niche cell types can be derived that specify both proliferative and quiescent niches for leukaemia.

The role of CDH2 in directing stem-cell fate, tumor-microenvironment interactions and chemo-resistance has been implicated in a wide variety of solid tumors and certain hematological malignancies like CML(*54, 55*). However, clinically relevant data on the role of CDH2 in microenvironment-mediated cancer proliferation and therapy resistance in acute leukaemia have until now been scarce as has means of clinically targeting this via low toxic agents. In a range of patient-derived leukaemia cells CDH2 binding and signaling is essential in mediating contact of the leukaemic blasts with niche cells: CDH2 is upregulated upon heterologous cell contact of the leukaemic cells with iMSC; in contrast, CDH2 knock-down in either leukaemic blasts or iMSC disrupts this interaction, leads to down-regulation of key oncogenic pathways, e.g. JAK/STAT signaling, in the leukaemic blasts and to an impaired survival and proliferation of the leukaemic blasts.

We detect that CDH2 is crucial in mediating microenvironment-mediated therapy resistance. Heterologous contact of the leukaemic blasts in these co-culture conditions induced expression of CDH2 and knock-down of CDH2 in MSC increased the sensitivity of patient-derived blasts by 3 fold. Interestingly, iMSC and vasculature niche-type iANG cells supported leukaemia survival under treatment in different ways. Dexamethasone treatment of patient-derived ALL cells on iANG primarily led to the selection and survival of blasts in G0 similar to what has been previously described. On iMSC, while there was still the emergence of a small resting blast cell population under treatment pressure with dexamethasone, the majority of surviving blasts continued to cycle. These cycling blasts expressed even higher levels of CDH2. Therefore, although a key feature of treatment resistance remains dormancy, not all treatment resistant cells are dormant and other mechanisms warrant further attention.

Ultimately, we confirmed that clinically relevant concentrations of the pentapeptide ADH- 1(*24*), a functional antagonist of CDH2 showed high efficacy against a panel of 15 patient derived leukaemia samples. ADH-1 efficacy similar to that of dexamethasone alone was confirmed in a xenograft mouse model of a highly aggressive incurable leukaemia. In this model, combination treatment with ADH1 and dexamethasone was more efficient than dexamethasone alone and the addition of ADH-1 did not increase toxicity. ADH-1 has been explored as an antiangiogenic drug in early phase clinical trials in solid tumors and published data indicate a tolerable clinical toxicity profile(*23–25, 39*) . ADH-1 may, therefore, be a candidate for clinical repurposing or a good starting poin t for a drug discovery program to meaningfully target the niche in blood cancers.

Using the complex hematopoietic bone marrow niche as a paradigm our proof-of-concept preclinical platform provides a prototype which can be adapted to investigate malignant niches in a wide variety of hematological cancers. As an example of proof of confidence in application our findings highlight the role of N-cadherin signaling in microenvironment-mediated drug resistance in leukaemia. This provides a starting point for the development of safer and more efficacious therapies to clinically target the tumor microenvironment.

## MATERIALS AND METHODS

### Study design

The aim of this study was to use leukaemia as a paradigm and detect a way to clinically target microenvironment mediated treatment resistance. In order to do this the study design had two goals: 1. To develop a tractable human cell based ex vivo BM milieu that would facilitate niche-mediated survival and proliferation of patient-derived cancer cells 2. To reveal microenvironment-dependent leukaemia biology including proliferation, dormancy and treatment resistance. We developed synthetic human BM niche cell types from BM mesencymal stroma reprogrammed iPSC. In vitro, BM-iPSC were differentiated into mesenchymal stem cells, perivascular and endothelia – like cells. We conducted in vitro co-culture experiments that BM-iPSC-derived niche cells could support human BM-derived haematopoietic cells (3 donors) and patient-derived leukaemia samples (14 samples). We conducted gene expression profiling in niche primed blasts to assay niche mediated changes in adherens junction molecules (2 samples) and confirmed upregulation of CDH2 via qRT-PCR in 7 samples (6 diagnostic and 1 relapse). Consequent functional validation experiments included CDH2 knockdown in leukaemia cells (4 cell lines) and patient-derived leukaemia cell co-culture on iMSC^CDH2-^ (3 diagnostic, 1 relapse sample). To validate if CDH2 could be therapeutically targeted in the clinics we performed in vitro drug dose response assays with CDH2-antagonist ADH1 (single drug assay: 15 diagnostic, 1 relapse samples, combination: 3 samples). Unless stated all experiments were conducted with a minimum of two independent experimental repeats, n=3. All graphical plots show standard deviation as error bars. All other imaging or flow cytometry data show a representative example of the total number of experiments

### Patient samples

Patient-derived leukaemia blasts were obtained from the Newcastle Biobank (REC reference number 07/H0906/109+5). Samples obtained from UCL were made under Research Ethics Committee reference 14/EM/0134.

### Mouse xenograft studies

All mouse studies were carried out in accordance with UK Animals (Scientific Procedures) Act, 1986 under project license P74687DB5 following approval of Newcastle University animal ethical review body (AWERB). Mice were housed in specific pathogen free conditions in individually ventilated cages with sterile bedding, water and diet (Irradiated No. 3 breeding diet, SDS). Mice were checked daily to ensure good health. All procedures were performed in a laminar flow hood except bioluminescent imaging (BLI).

NSG mice (NOD.Cg-Prkdc^scid^ Il2rg ^tm1Wjl^/SzJ) aged between 12 and 16 weeks, both male and female, from in-house colonies were used for transplantions. Mice were checked daily, weighed and examined at least once weekly during studies to ensure good health.

### Statistics

A two-way analysis of variance, multi comparison with Tukey test was used to compare in vivo efficacy group total flux measurements from bioluminescent imaging. One way analysis of variance was used to compare engraftment of BM and spleen and, spleen weight for *in vivo* efficacy treatments. All statistical tests were performed using GraphPad Prism 6.

## SUPPLEMENTARY MATERIALS AND METHODS

### BM-iPSC reprogramming and culture

iPSC reprogramming was performed on mesenchymal stroma cells isolated from bone marrow of hip replacement surgeries. Low passage stroma cells seeded at a density of 18,500 cells/cm2 were transfected using Simplicon™ RNA Reprogramming Kit, OKSG (Merck Millipore) following pre-treatment with B18R protein. Following puromycin selection over an 8 day period, cells were subjected to 10µg/ml of bFGF (GIBCO), 1µl/ml of Human iPS Reprogramming Boost Supplement II, 1000x (Merck Millipore) and mouse embryonic fibroblast conditioned media (R&D Systems). iPSC colonies appeared between Day 28-30 post transfection and were picked under a stem cell microdissection cabinet for subsequent cultures. iPSC cultures were maintained on Vitronectin XF™(Stemcell Technologies) coated plates in TeSR™2 media (Stemcell Technologies). BM-iPSC were subsequently differentiated to generate iMSC and iANG cells.

### BM-iPSC differentiation

BM-iPSC lines were differentiated into mesenchymal stem cells, endothelia and perivascular cells through an intermediate early mesoderm route using protocols adapted from existing studies(*56*). Briefly, mesoderm induction was carried in Mesoderm Induction Media (Stemcell Technologies) for 72 hours following which the cells were subjected to either mesenchymal or vascular specification steps. Messenchymal differentiation was achieved by treating the early mesoderm cells with Low-glucose Dolbecco’s Modified Eagle’s Medium (SIGMA), 20% Heat Inactivated Foetal Bovine Serum (GIBCO) and 10µg/ml of bFGF (GIBCO). Vascular specification was achieved by treating cells with Mesoderm Induction Media, 1µM Human Recombination VEGF-165 (Stemcell Technologies) and 1µM SB431542 (Stemcell Technologies) for 12 days following which CD31+ cells were sorted using flow cytometry for the purposes of characterisation. All cells following vascular specification were maintained in Microvascular Endothelial Cell Growth Medium (Sigma-Aldrich) for subsequent co-cultures.

### Ex vivo co-cultures

Patient derived leukaemia samples were seeded on iMSC or iANG cultures at a seeding density of 0.5-1 million cells/ml in SFEMII media (Stemcell Technologies) using protocols adapted from existing studies(*16, 53*).

### Cell generational tracing

10mM CellTrace^TM^ Violet (Life Technologies), Excitation/Emission: 405nm/450nm was used to stain patient derived leukaemia cells at a cell density of 1 million/ml in 1X phosphate buffered saline for a total of 20 minutes at 37C, 5% CO2 following which excess stain was removed and cells were immediately put into co-culture in SFEMII media for subsequent cell fate tracking and/or sorting using flow cytometry.

### Cell cycle and G0 analysis

Following co-culture cells were harvested as per existing protocols(*16*) and subsequently stained with 10µg/ml of Hoechst33342 (Sigma-aldrich), Excitation/Emission: 350nm/450nm for 45 minutes at 37C, 5% CO2 at a cell density of 1 million/ml in SFEMII media. Following this 5µl of 100µg/ml Pyronin Y (Sigma-aldrich), Excitation/Emission: 480nm/575nm was added to each 1 million/ml sample and stained for a further 15 minutes in the same conditions. Samples were then transferred onto ice and analysed by flow cytometry.

### FISH

5 million cells were pelleted through centrifugation for 3 minutes at 1,200 rpm, supernatant was subsequently discarded and 10ml 0.075M potassium chloride (pre-heated to 37 degrees) was added dropwise whilst mixing on a vortex. Samples were Incubated for 10 minutes at 37°C and further centrifuged at 1,200 rpm for 5 minutes. Supernatant was discarded and pellet vortexed. Following this 1ml of fresh fixative (3:1 methanol: acetic acid) was added dropwise with continuous vortexing which was then topped up to 5ml. Following another centrifugation step 1ml of fresh fixative was added for subsequent hybridisation procedure.

Briefly, 0.2ul of FISH probes (Dakocytomation TCF3 FISH DNA probe split signal, Agilent for E2A/HLF samples OR RP11-773I18 fluorescently labelled BAC probes(*57*) to detect RUNX1 amplification in iAMP21 samples) were mixed with 2.8ul hybridisation buffer (Cytocell, New York, USA) and denatured at 75 °C for five minutes followed by hybridisation at 37 °C overnight. Coverslips were removed in 2x SSC and slides washed in 0.02% SSC with 0.003% NP40 at 72 °C for two minutes followed incubation in 0.1% SSC at room temperature for two minutes. Slides were mounted with 10 ul DAPI (Vector laboratories, California, USA). Scoring was performed on an automated Olympus BX-61 florescence microscope with a ×100 oil objective using CytoVision 7.2 software (Leica Microsystems, Newcastle-upon-Tyne, UK). Where possible, more than 100 nuclei were scored for each FISH test by two independent analysts. A cut-off threshold of >5% was established by counting the number of abnormal (false positive) signals generated when probes were hybridised to normal cells.

### Immunofluorescent Staining

Immunofluorescent staining was conducted on Vitronectin ^TM^ coated EZ chamber slides. B M-i P S C co l o n ies w e re fixed using 4% formaldehyde solution for 20 minutes at room temperature. The cells were then washed twice with 1x PBS for 10 minutes. For nuclear staining, cells were permeabilised using 0.1% triton X-100/1xPBS for 10 minutes at room temperature, then washed twice with 1x PBS for 10 minutes. Blocking solution (4% normal goat serum) was added for 30 minutes at room temperature. The cells were then incubated with the primary antibody (table S4) using a 1:250 dilution overnight and then washed three times with 1x PBS for 10 minutes. Subsequently cells were incubated with the secondary antibody in 1:500 dilution for 60 minutes at room temperature before being washed three times with 1xPBS for 10 minutes. N u c l e a r co u n te rsta i n DAPI was added to each well in 1:500 dilution and incubated at room temperature for 10 minutes. Finally, the coverslip was mounted onto a slide using gold antifade reagent and slides were visualized using the Nikon A1 confocal fluorescent microscope.

### Alkaline Phosphatase detection

BM-iPSC were cultured for a minimumof 5 days, when alkaline phosphatase (AP) signal is optimal. On day 6, the cells were washed three times in PBS for 10 minutes and fixed using 4% paraformaldehyde for 2 minutes. The cells were washed with 1X rinse buffer (TBST-20mM Tris-HCL, pH 7.4, 0.15 NaCl, 0.05% Tween-20). Alkaline phosphatase staining solution was prepared fresh by mixing Fast Red Violet (FRV) with Naphthol AS-BI phosphate solution and sterilised distilled water in a 2:1:1 ratio and added to cover the base of the well for a 15 minute incubation in the dark. Subsequently cells were washed with 1x PBS and stored in PBS till analysis. Positively stained iPSC colonies could be seen by eye, a microscope was used to visualise the colonies in greater detail.

### Sequencing and analysis

#### mRNA-sequencing and analysis

Sequencing libraries were prepared using the TruSeq Stranded mRNA Sample Preparation Kit [Illumina] following manufacturer’s instructions. Pooled libraries were sequenced at 40 Million (2 x 75 bp) reads per sample using a NextSeq 500 and High Output Kit (150 cycles) [Illumina]. The quality of sequenced reads was assessed using FastQC(*58*), which suggests high quality data with all reads have Phread score > 30 across all bases. For each sample, transcript abundance was quantified from raw reads with Salmon (version 0.8.2)(*59*) using the reference human transcriptome (hg38) defined by GENCODE release 27. An R package Tximport (version 1.6.0)(*60*) was used to estimate gene-level abundance from Salmon’s transcript-level counts. DESeq2 (version 1.18.1)(*61*) was used to generate gene-level normalized counts and to perform differential expression analysis.

#### Whole-exome sequencing data analysis

Sequencing libraries were prepared using the Nextera Rapid Capture Exome Kit [Illumina] following manufacturer’s instructions. Pooled libraries were sequenced at >90X coverage (2 x 75 bp) per sample using a NextSeq 500 and High Output Kit (150 cycles) [Illumina].Raw reads were aligned to human reference genome (hg19) using Burrows-Wheeler Aligner (BWA) 0.7.12(*62*) and were processed using the Genome Analysis Toolkit (GATK, v3.8) best practices recommended workflow for variant discovery analysis(*63–65*). MuTect (v1.1.7) and MuTect2(*66*) were used to identify somatic variants (SNPs and INDELs) in the iMSC and iANG primed patient derived blasts that were not present in the blasts prior to co-culture. Variants were annotated using Ensembl Variant Effect Predictor (VEP, version 90)(*67*). Circos plots of exonic mutations with allele frequency > 25% were generated using Circos(*68*).

### Quantitative RT-PCR

RNA was extracted using Qiagen RNEasy Micro Procedure as per manufacturer’s protocol following an on column DNAse removal step. RevertAidTMH Minus First Strand cDNA Synthesis Kit (ThermoFisher Scientific) was used to synthesise cDNA. 500ng RNA was collected and added to RNase/DEPC free water to a final volume of 11μl. 1µl (dN)6 (200mg/l) random hexamers was added, mixed gently by inverting the vial and briefly centrifuged. Using a GeneAmp PCR system 2700 the sample was incubated at 65°C for 5 minutes, after which the sample was immediately placed on ice, 8µl of the master mix (5X Reaction Buffer, 20U/µl RNAse Inhibitor, 10mM dNTP and 100 U/µl RevertAid H Minus MMLV RT) was added, the samples were vortexed and briefly centrifuged. The samples were placed back in the PCR machine to incubate at 25°C for 10 minutes, 42°C for 60 minutes and 75°C for 10 minutes to terminate the reaction.

Primers (table S4) were reconstituted in RNase/DNase free water to a working solution of 10µM. The PCR master mix (reverse and forward primer, SyBr-Green master mix and RNAse free water) was mixed well by gently pipetting the solution, 8μl/well was added to a 384-well PCR plate, 2μl cDNA was then added to each well to a total of 10µl/well. The plate was sealed and centrifuged for 1 minute at 1000RPM and placed in an applied Biosystems 7900HT Sequence Detection System and ran 40 cycles. This included a denaturation step at 95°C, an annealing step at 60°C and an elongation step at 90°C. For RT-PCR arrays cDNA was synthesised using RT2 First Strand Kit (Qiagen) and subsequent PCR step were performed same as above but using RT2 SYBR Green ROX qPCR Mastermix (Qiagen) and PAHS-086ZE-4 - RT² Profiler™ arrays (Qiagen) as per manufacturers protocol.

### Functional in vitro knockdown

#### Leukaemia cell lines

ALL and AML cell lines SEM, HAL-01, PreB 697 and KASUMI were authenticated by short tandem repeat profiling by NewGene Ltd (Newcastle University, UK). Cell lines were confirmed free from mycoplasma infection at regular intervals using a MycoAlert kit (Lonza, Slough, UK). Cells lines were routinely maintained in RPMI-1640 medium supplemented with 20% FBS and L-Glutamine.

In order to engineer a doxycycline conditional RNAi approach oligos (shRNA guide strand) designed against CDH2 were cloned into a pL40C.T3.dTomato.miRN.PGK.Venus.IRES.rtTA-V10.WPRE backbone as per published protocols(*42, 69*). Following bacterial transformation and plasmid amplification DNA sequence was confirmed through sanger sequencing. Plasmid DNA was obtained using Endofree® Plasmid Kit (Qiagen) and introduced into 293T cells for lentivirus production. The 293T cells were grown in 150 mm tissue culture dishes at a concentration of 3 x 106 cells in 30 ml DMEM media the day prior to the co-transfection. On the following day, 45 µg packaging plasmid pCMVΔR8.91, 15 µg envelope plasmid pMD2.G, and 60 µg shRNA expression vector were co-transfected using calcium phosphate precipitation method. The cells were incubated for 72 hours and the recombinant pseudotyped lentivirus-containing supernatant was collected for subsequent concentration of the thus engineered lentivirus. Lentiviral transduction was performed on leukaemia cell lines as described previously and transduced (VFP+ve), doxycycline induced cells (dTomato+ve) were selected by flow cytometry.

#### iMSC

iMSC were transduced with sc-29403-V N-cadherin shRNA lentiviral particles (Santacruz Biotechnology) as per manufacturer’s protocols and stable cell lines expressing the shRNA were isolated using puromycin selection.

#### Drug dose response

Single agent and combinatorial drug dose response assays were set up as previously described(*16, 53*). Briefly, patient-derived leukaemia cells were seeded at 0.5-1 million/ml density onto iMSC or iANG cells. Clinically relevant concentration of different treatment compounds were added 24 hours later and cells were harvested for manual counting after a 5 day period.

#### Mouse in vivo studies

Mice were housed in specific pathogen free conditions in individually ventilated cages with sterile bedding, water and diet (Irradiated No. 3 breeding diet, SDS). Mice were checked daily to ensure good health. All procedures were performed in a laminar flow hood except bioluminescent imaging (BLI).

##### Leukaemia PDX cell production

NSG mice were injected with 1x10E^4^ -1x10^6^ cells in 20-30µl/mouse in RPMI1640 (SIGMA), 10% FBS (Sigma) intrafemorally (i.f.) directly into the femur bone marrow. During the procedure mice were anesthetised by isoflurane inhalation and provided with analgesia (Carprofen, 5mg/kg subcutaneously with 29G needle). Mice were humanely killed at a time point prior to adverse health effects as determined by previous studies and the presence of an enlarged spleen visible through the skin of the abdomen. Any mice that displayed symptoms of leukaemia such as weight loss, anaemia, and hypotonia were immediately humanely killed. PDX cells were harvested from the spleen via cell disruption though a cell strainer (40µm, SLS Ltd.), washed twice in sterile PBS and stored frozen in 10%DMSO;90%FBS (Sigma).

##### Teratoma studies

5 NSG mice (group size determined from previous studies) per i-niche sample were injected with 5x10^5^ cells 1:1 in Matrigel (Standard formulation, Corning Inc.) subcutaneously in a volume of 100µl per mouse on the flank with a 29G needle.

Tetratoma formation was assessed and measured using calipers at least once weekly. Mice were humanely killed when tumours reached 1.5cm diameter in any direction. Masses were dissected and fixed in formalin for H&E staining by standard methods.

##### Engraftment of i-niche cultured PDX cells

3x10^5^ L707D PDX Luc^+^ GFP^+^ cells following culture with either iMSC or iANG were injected i.f. into 3 mice/niche. Engraftment was assessed via BLI (see below) and spleen size.

##### ADH1/Dex in vivo efficacy study

A dose escalation toxicity test was performed in 2 female and 2 male NSG mice to determine a tolerated dose and schedule. 20 NSG mice were injected i.f with 1x10^4^ L707D PDX Luc^+^ GFP^+^ 1x10^5^ Luc^+^ GFP^+^ cells in 20µl media/mouse as described above. Mice injected with this PDX have an event free survival of 4-5 weeks. The study was designed to end 28 days after transplant to maximise the number of PDX cells harvested, to minimise any mouse ill health and to compare treatment effect by comparison of tissue engraftment. Six days after injection, mice were randomised into 4 treatment groups. Mice were housed in a least two cages per treatment group to minimise cage effects. Five mice per group was calculated to be the minimum number to identify a significant difference in BLI total flux between the groups after 3 rounds of dosing. Treatments were administered via intraperitoneal injection using a 29G needle and saline (0.9% w/v) vehicle in a volume of 5 µl/g mouse weight. ADH1 (Adooq Bioscience) was dissolved in saline just before injection. Dexamethasone sulphate solution was diluted in saline and combined with ADH1 for a single injection. Groups were given either saline (CV), 3mg/kg dexamethasone (Dex), 200mg/kg ADH1 or Dex/ADH1 combined (3mg and 200mg/kg respectively), 1x daily, 5x weekly for 3 weeks. Engraftment was assessed via bioluminescent imaging (IVIS Spectrum, Caliper with Living Image Software). For imaging, mice were injected with 150mg/kg d-luciferin interperitoneally (In vivo Glo, Promega) and anaesthetised with isoflurane. Mice were humanely killed, and spleen cells harvested as described above. A portion of spleen was fixed in formalin for immunohistochemistry. Muscle was removed from all leg and hip bones and bone marrow (BM) cells were isolated by crushing the bones in PBS in a pestle and mortar and washing the bone fragments with PBS.

##### Engraftment assessment of mouse spleen and BM

Isolated cells were counted and suspended in 0.05%BSA (Roche) in PSB. Cells were stained with mouse CD45 PeCy7 and human CD45 FITC (BD Biosciences) following suppliers’ instructions and analysed by flow cytomentry (Attune, Thermo) Fixed tissues from the efficacy study mice were processed for immunochistochemistry by Cellullar Pathology, RVI Newcastle Hospitals NHS trust using standard methods as follows. Briefly, bones were decalcified using EDTA. Tissues were infiltrated with and embedded in parrafin wax. Sections on slides were staining with either, hematoxylin and eosin (H&E) or, human CD19 antibody using a Ventana BenchMark Ultra (Ventana, Roche), and Universal DAB Detection kit (Ultraview) to produce a brown chromogen at the site of human CD19. Slides were scanned using an Aperio ScanScope (Leica) and images analysised using Leica eSlide manager software.

## Supplementary figures

**S 1.**
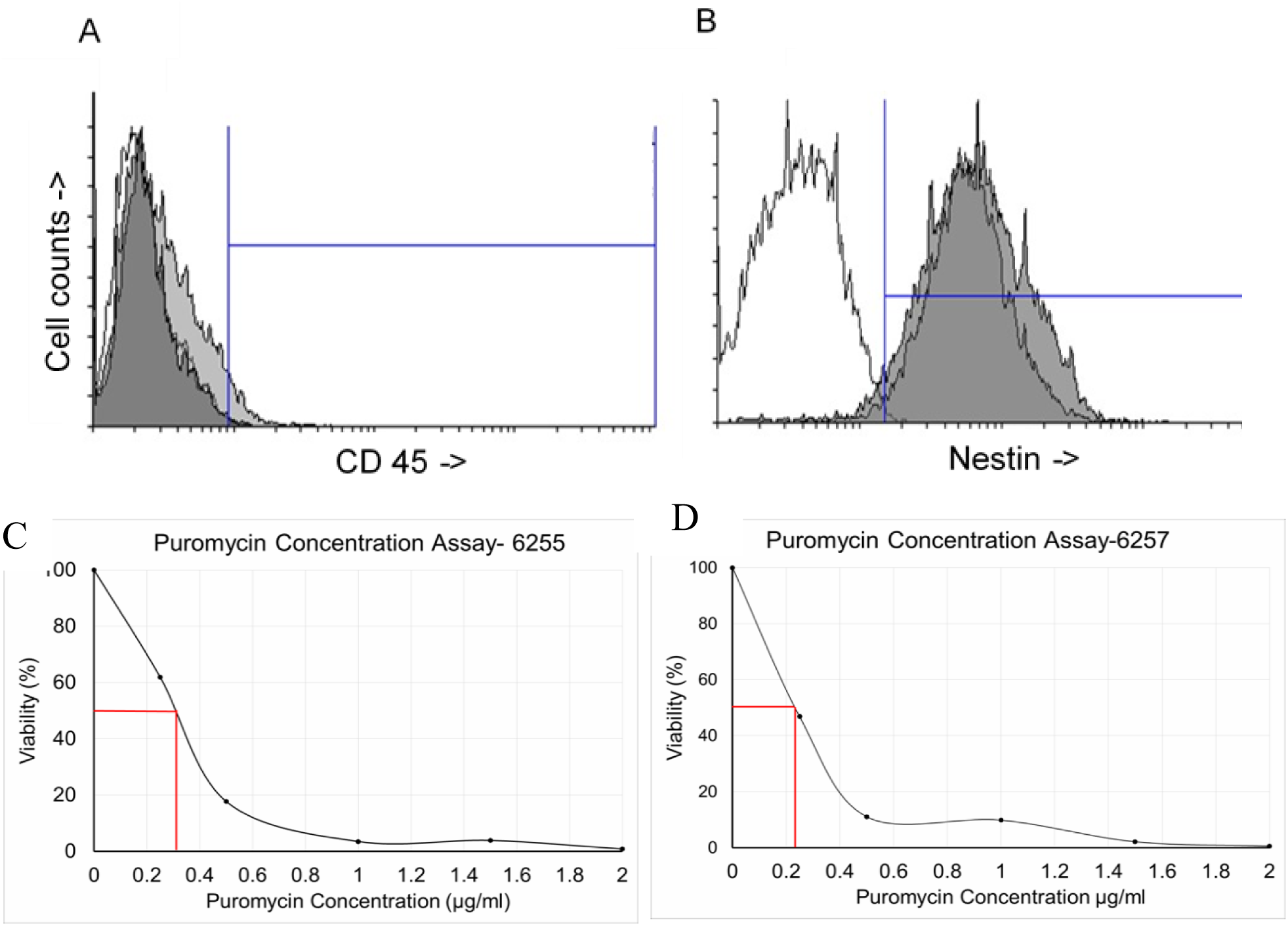
CD45 and Nestin expression in primary human mesenchymal stroma cells. Flow cytometry analysis shows unstained control cells (white) negative for both CD45 (A) and Nestin (B). Both mesenchymal stroma samples 6255 (light grey) and 6257 (dark grey) both are negative for CD45 expression but positive for Nestin expression. C-D. Puromycin Optimisation kills curves for primary human bone marrow mesenchymal stroma cells 6255(C) and 6257(D) with IC50 indicated by red line (0.35 μg/ml and 0.38 μg/ml respectively).

**S2.**
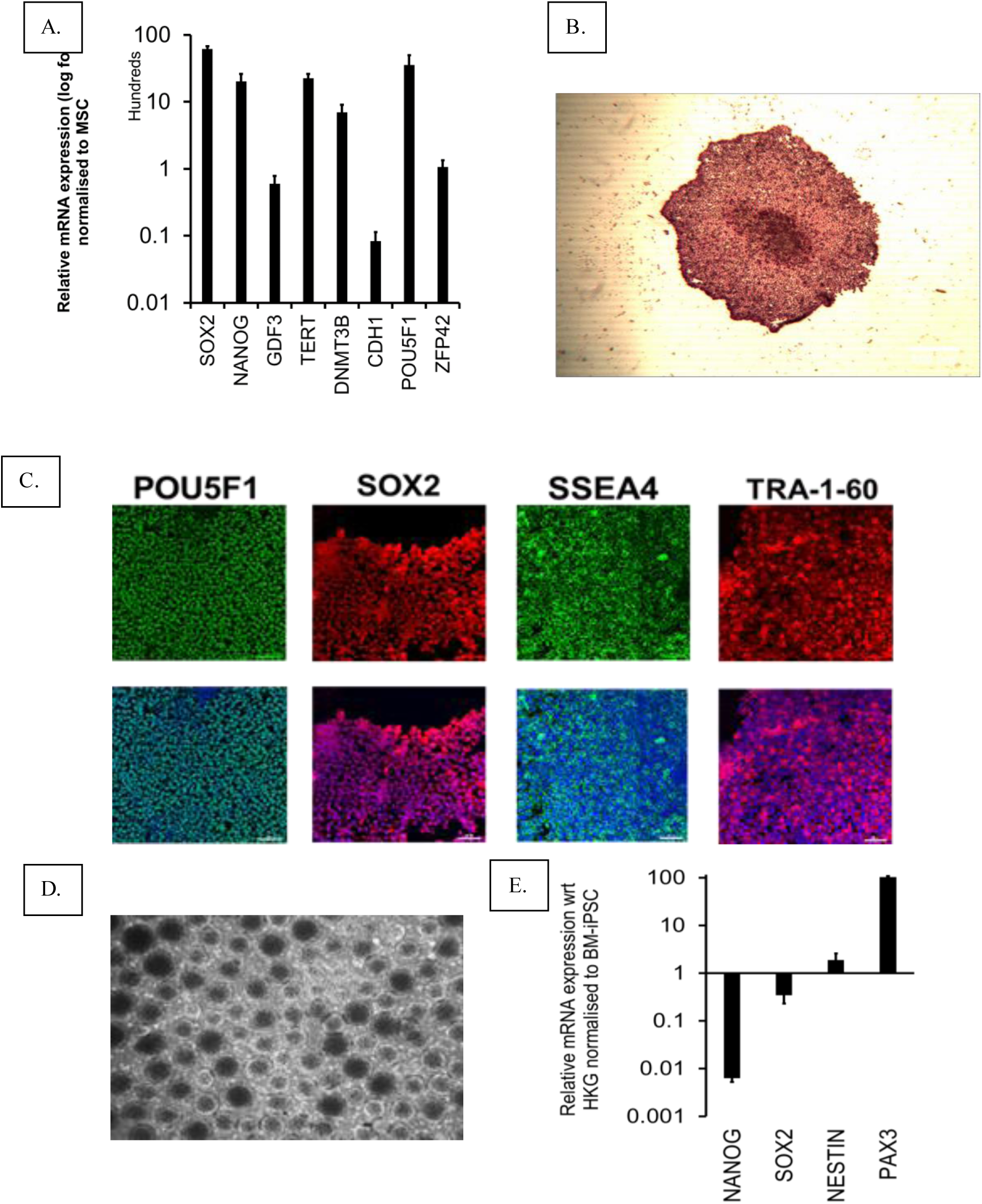
A. Relative mRNA expression of pluripotent transcripts in BM-iPSC normalised against parental bone marrow stroma B. Image showing BM-iPSC stained for Alkaline phosphatase C. Nuclear POU5F1 and SOX2, cell surface SSEA4, TRA-1-60 staining in feeder-free and xeno-free cultures of BM-iPSC D. Optical image of in vitro embryoid bodies E. In vitro differentiation of BM-iPSC derived embryoid bodies down-regulates pluripotent transcripts and upregulates ectoderm and mesoderm genes Nestin and PAX3 respectively.

**S3.**
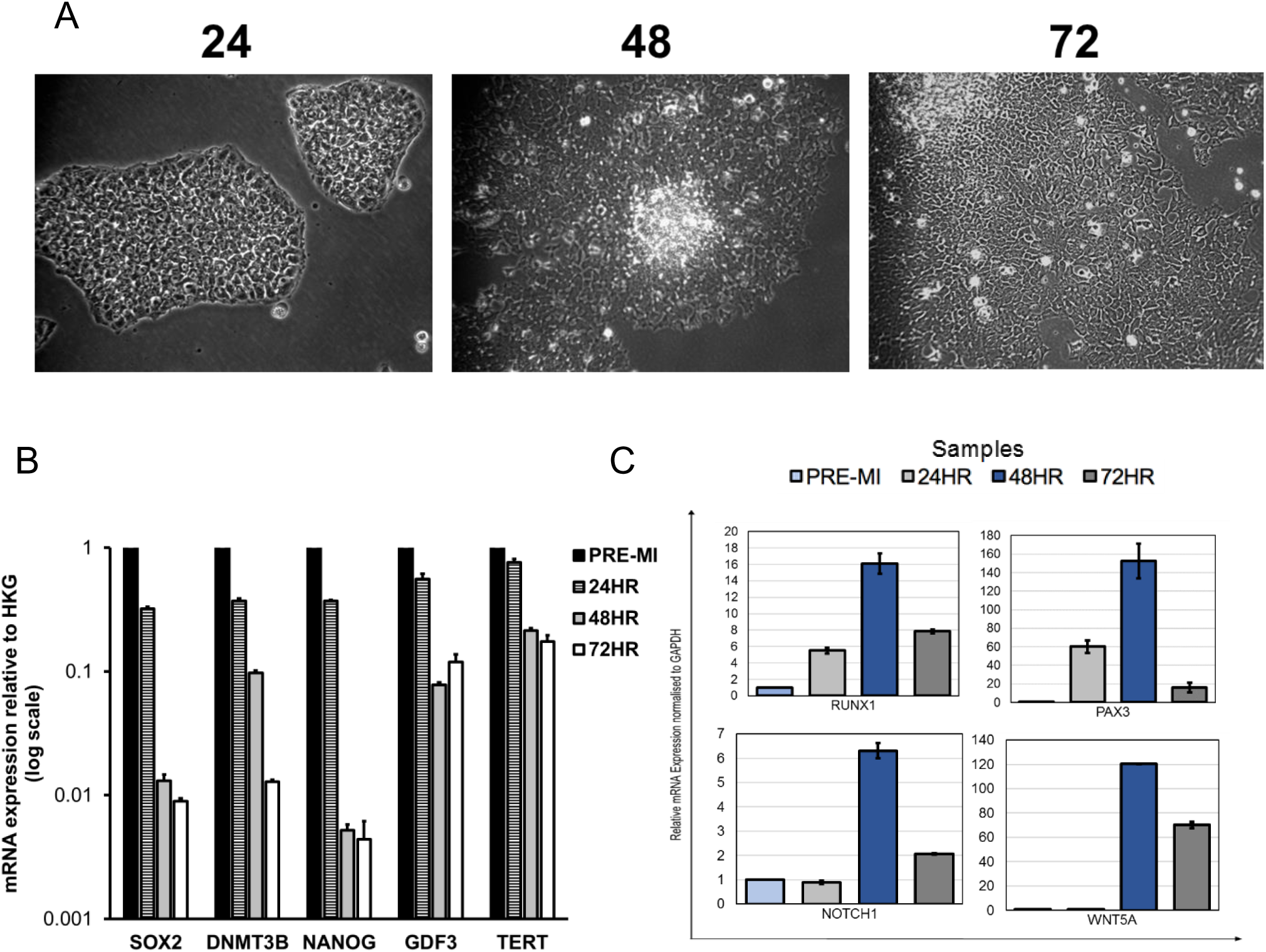
A. Photomicrograph of BM-iPSC differentiation into early mesoderm cells over 72 hours B. mRNA expression of pluripotent transcripts during mesodermal differentiation of BM-iPSC C. Relative mRNA expression of mesoderm genes RUNX1, PAX3 and NOTCH1 and WNT5A during mesodermal differentiation of BM-iPSC

**S4.**
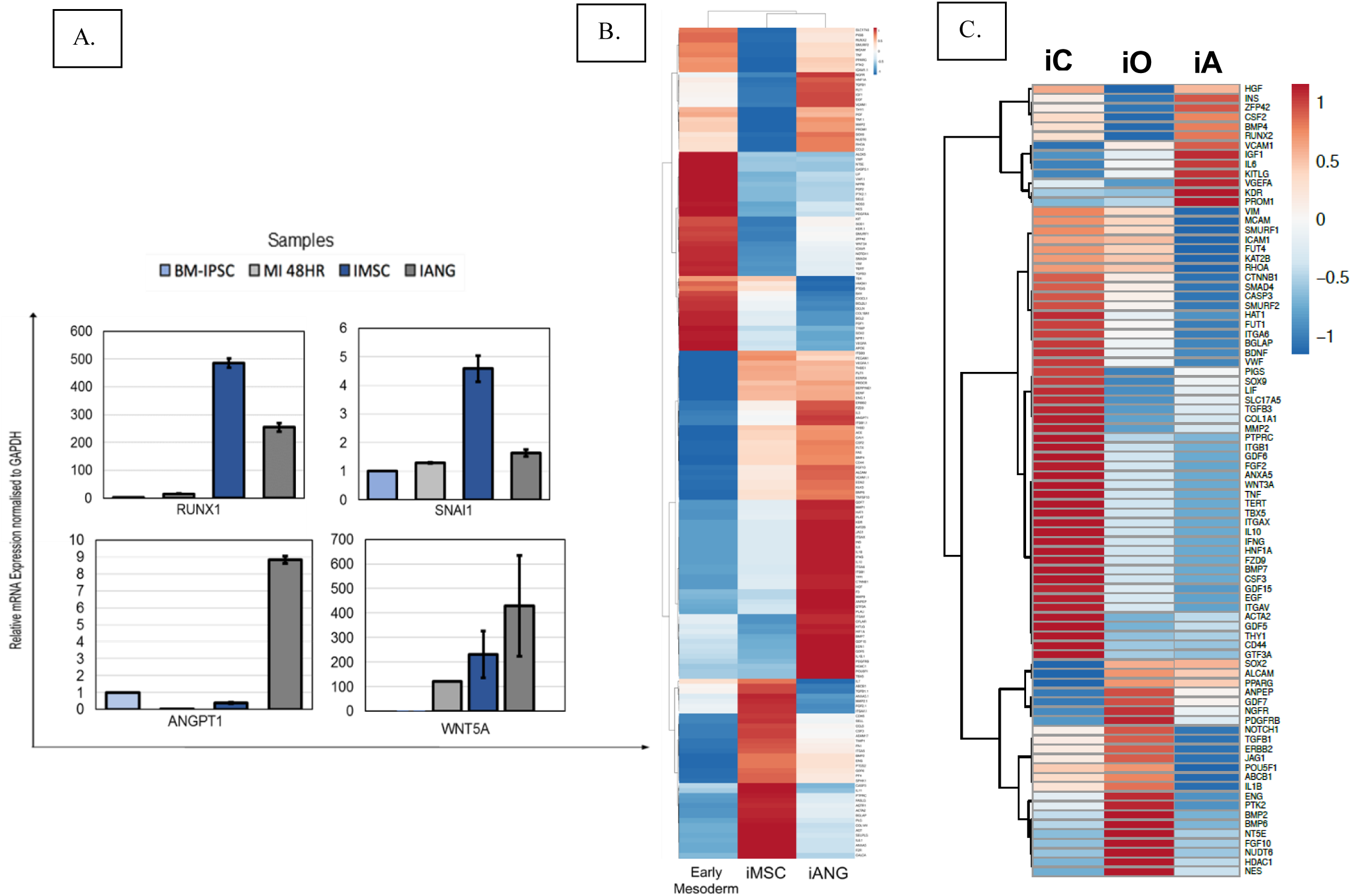
A. Relative mRNA expression of messenchymal genes RUNX1, SNAI1 and ANGPT1 and WNT5A during in early mesoderm (mesoderm induction, MI at 48 hrs), iMSC and iANG B. BM-iPSC derived early mesoderm, mesenchymal [iMSC] and vascular [iANG] cells demonstrate distinct transcriptomic profiles as evaluated by high throughput gene expression arrays C. iMSC differentiate into chondrocytes [iC], osteocytes [iO] and adipocytes [iA] with distinct gene expression profiles.

**S5.**
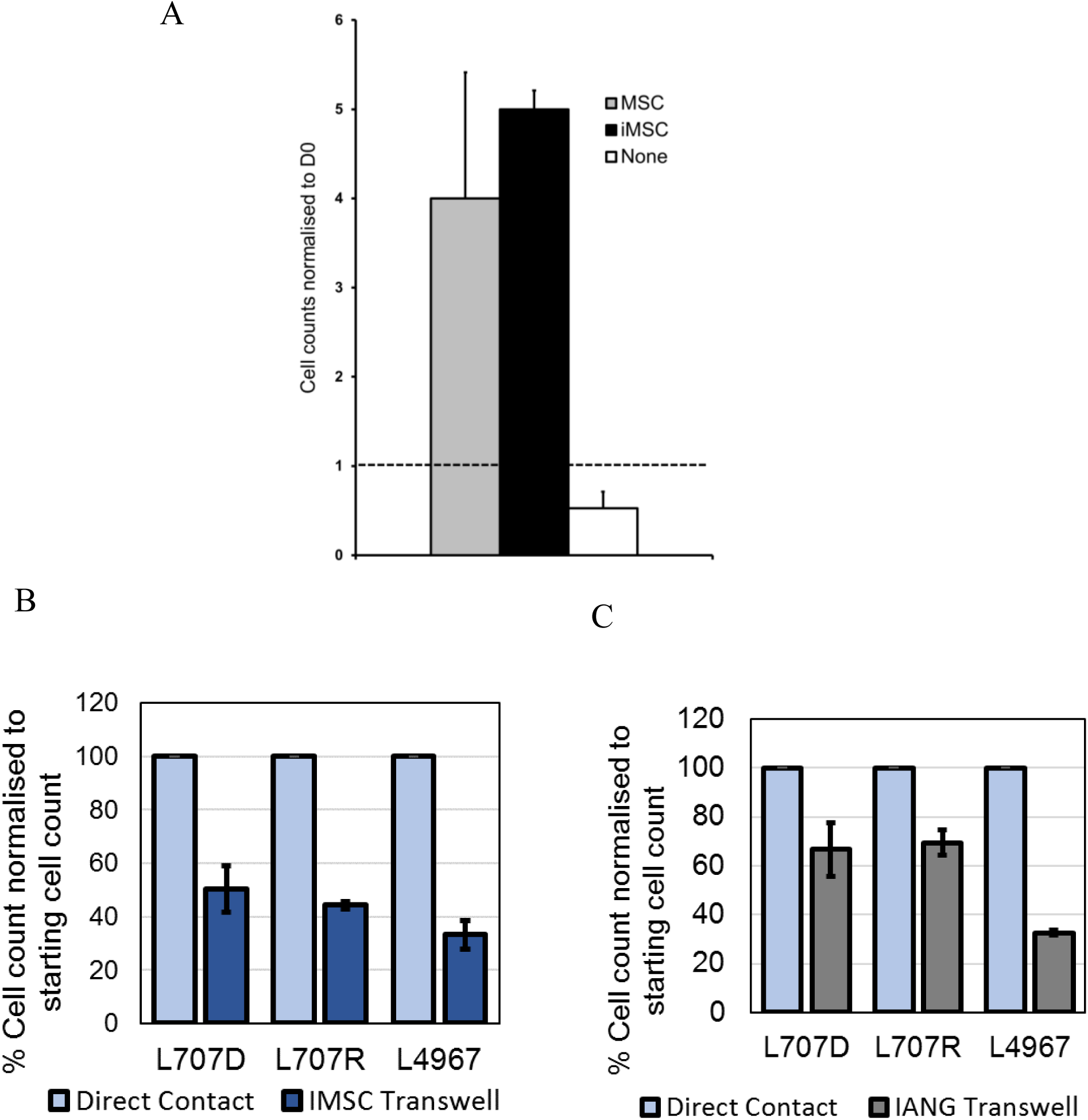
A. Cells counts of patient-derived acute lymphoblastic leukaemia cells on primary MSC, MSC, BM-iPSC derived MSC [iMSC] and in feeder-free suspension [None]. B-C. Cell counts of patient leukaemia ALL cells on direct contact and transwell conditioned media co-cultures with B. iMSC and C. iANG. L707D, L4967 = samples at diagnosis; L707R = matched relapse sample

**S6.**
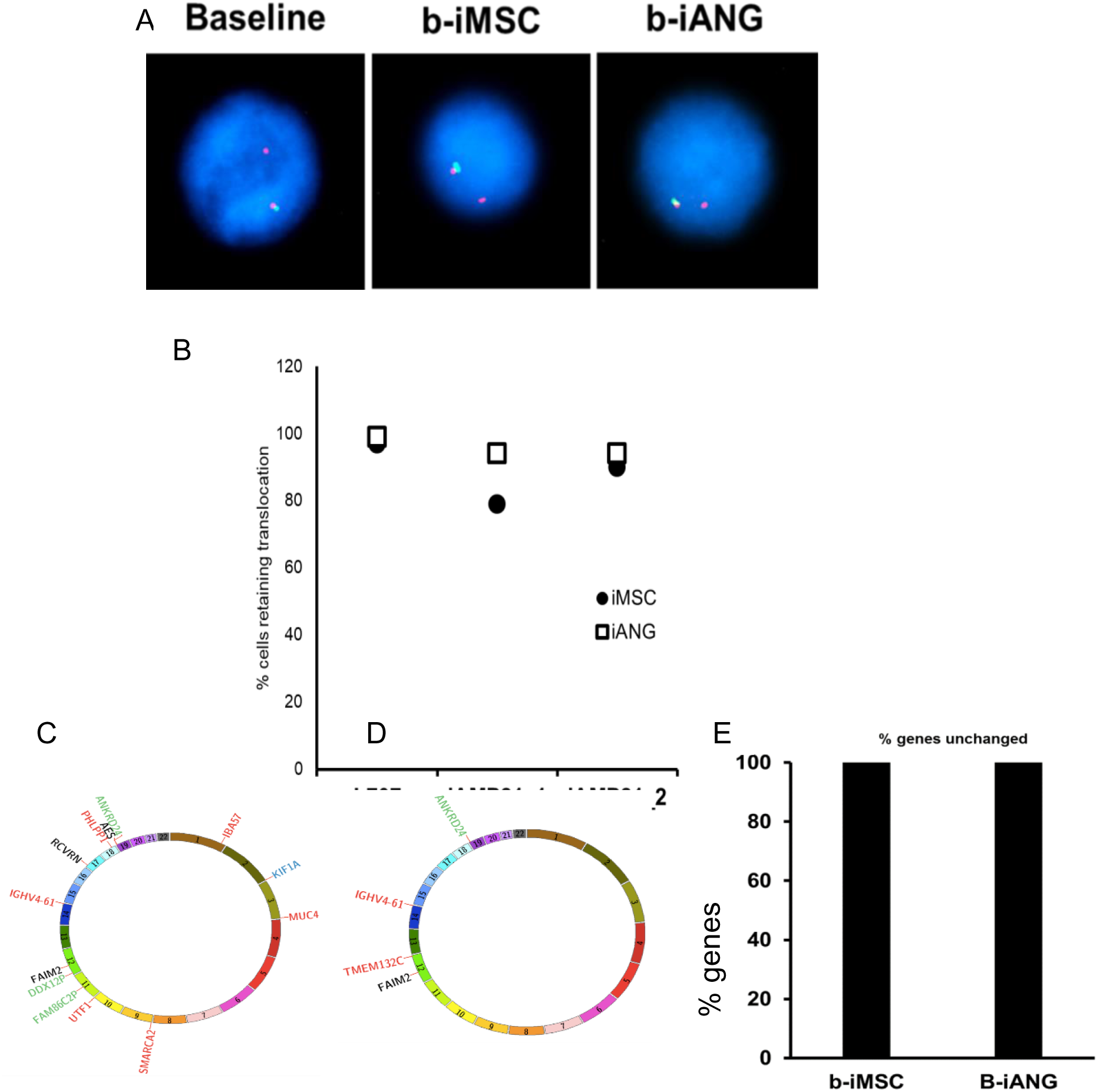
A. FISH images of E2A-HLF break-apart probe showing 1 breakpoint and 1 intact loci in baseline and retention of this staining in blasts primed with iMSC, b-iMSC and ANG, b-iANG B. Scoring data confirms retention of initial cytogenetic translocation in patient blasts following-niche co-culture. L707 = confirmation of E2A breakpoint; iAMP21 = confirmation of additional copies of RUNX1 which is a feature of iAMP21 (abnormal amplification of chromosome 21) samples. C-D. Circos plots showing whole exome changes in patient blasts following co-culture on C. iMSC and D. iANG. Over a period of 4 weeks. Green=silent:synonymous_variant/non_coding_transcript_exon_variant, black= UTR_variant, red = missense variant, blue = inframe deletion E. % exomes that are unchanged in patient blasts following i-niche co-culture.

**S7.**
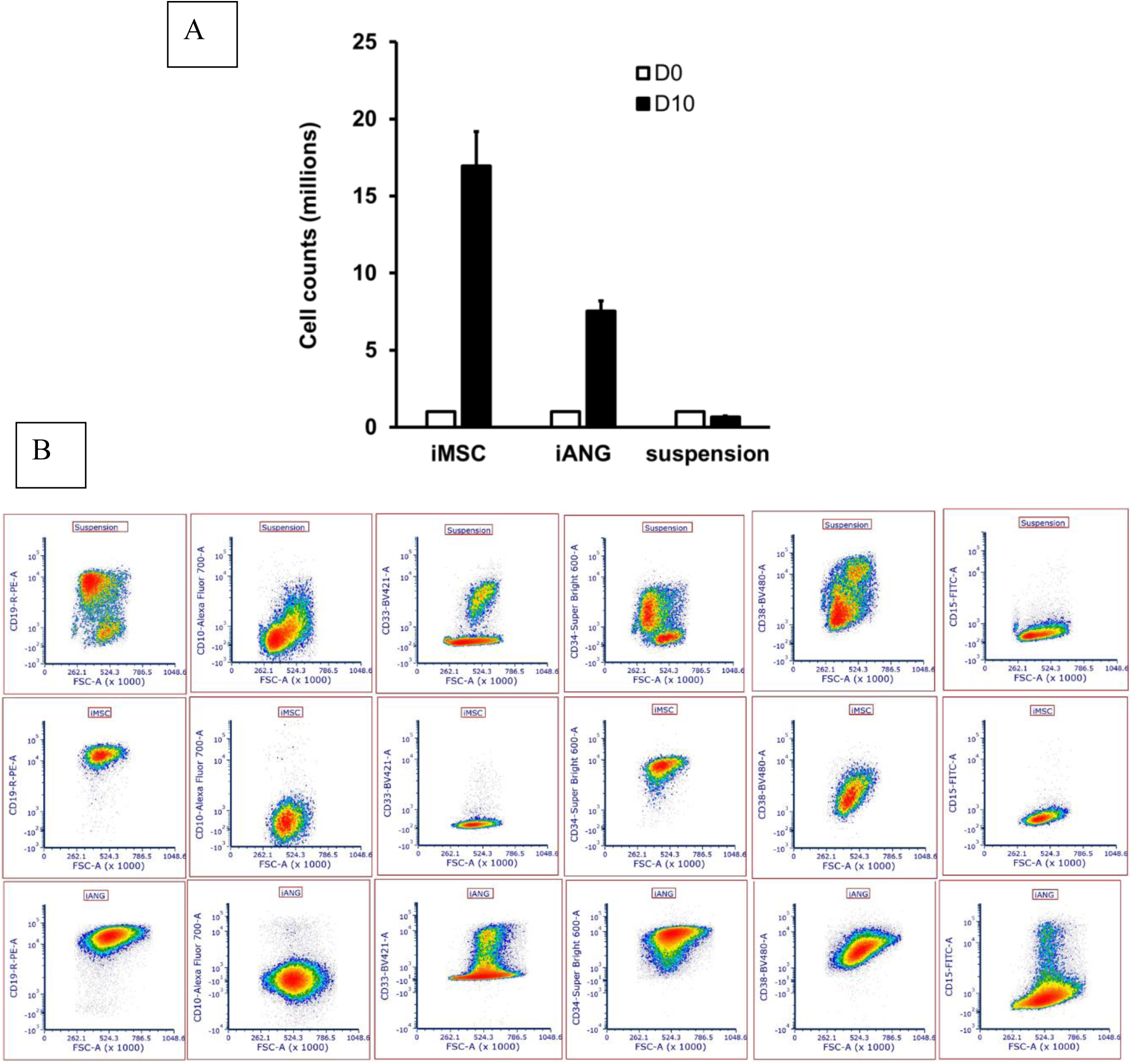
A Cell counts of blasts from a patient with biphenotypic MLLre leukaemia [MS40] on iMSC, iANG and in niche-free suspension cultures over 7 days. B. Immunophenotyping of MS40 blasts in niche-free suspension culture [top panel] and MS40 blasts primed by iMSC and iANG [middle, bottom panel] after 7 days

**S8.**
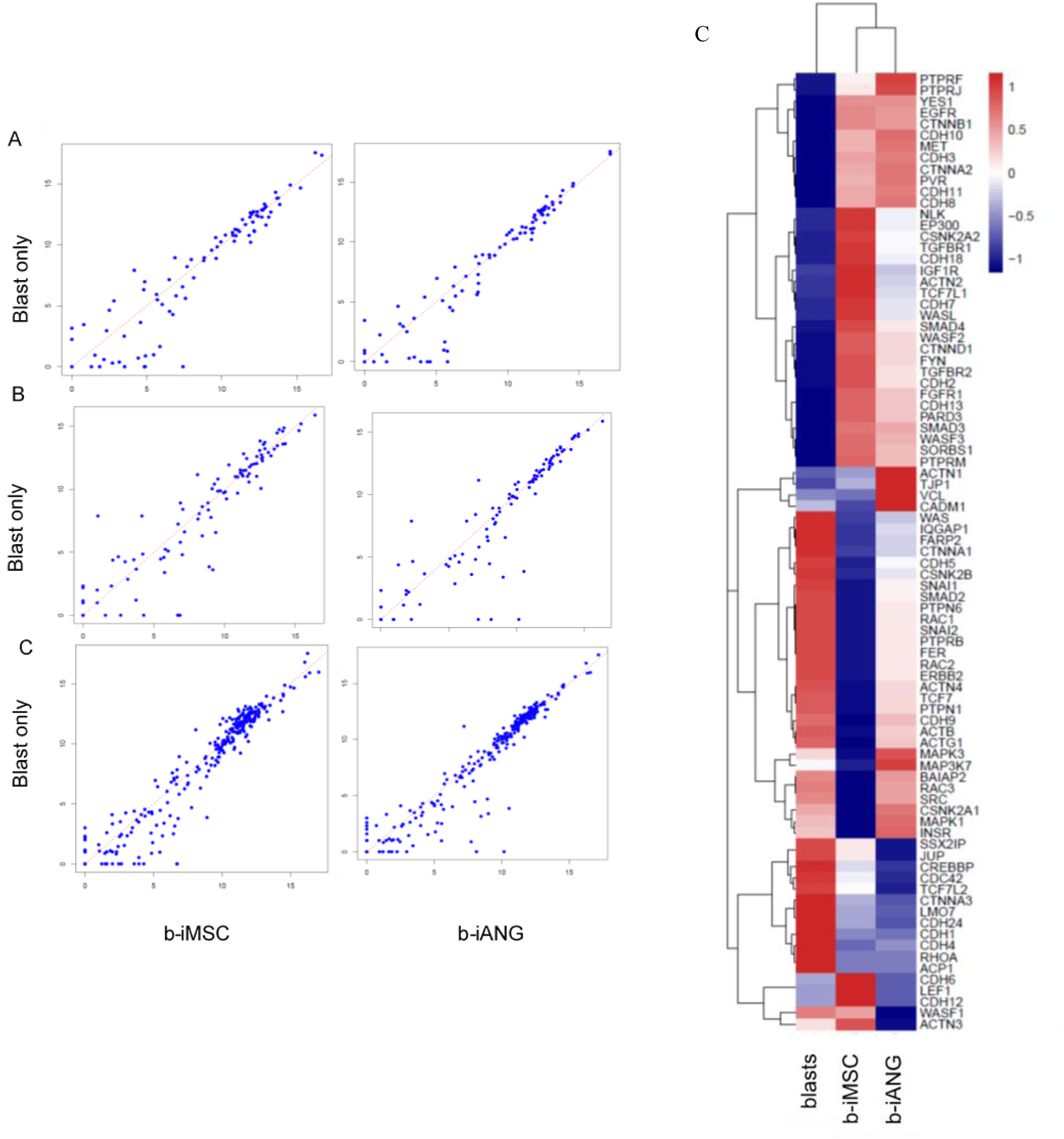
Scatter plots showing A. Adherens B. WNT and C. β-Catenin pathway genes in patient leukaemia blasts before (blasts only) and after i-niche co-culture for 7 days (b-iMSC, b-iANG) C. RNA Sequencing data showing adhesion molecules expression in patient leukaemia cells, sample L707D. Blasts before co-culture are compared to blasts following a 7-day co-culture on iMSC (b-iMSC) and iANG (b-iANG)

**S9.**
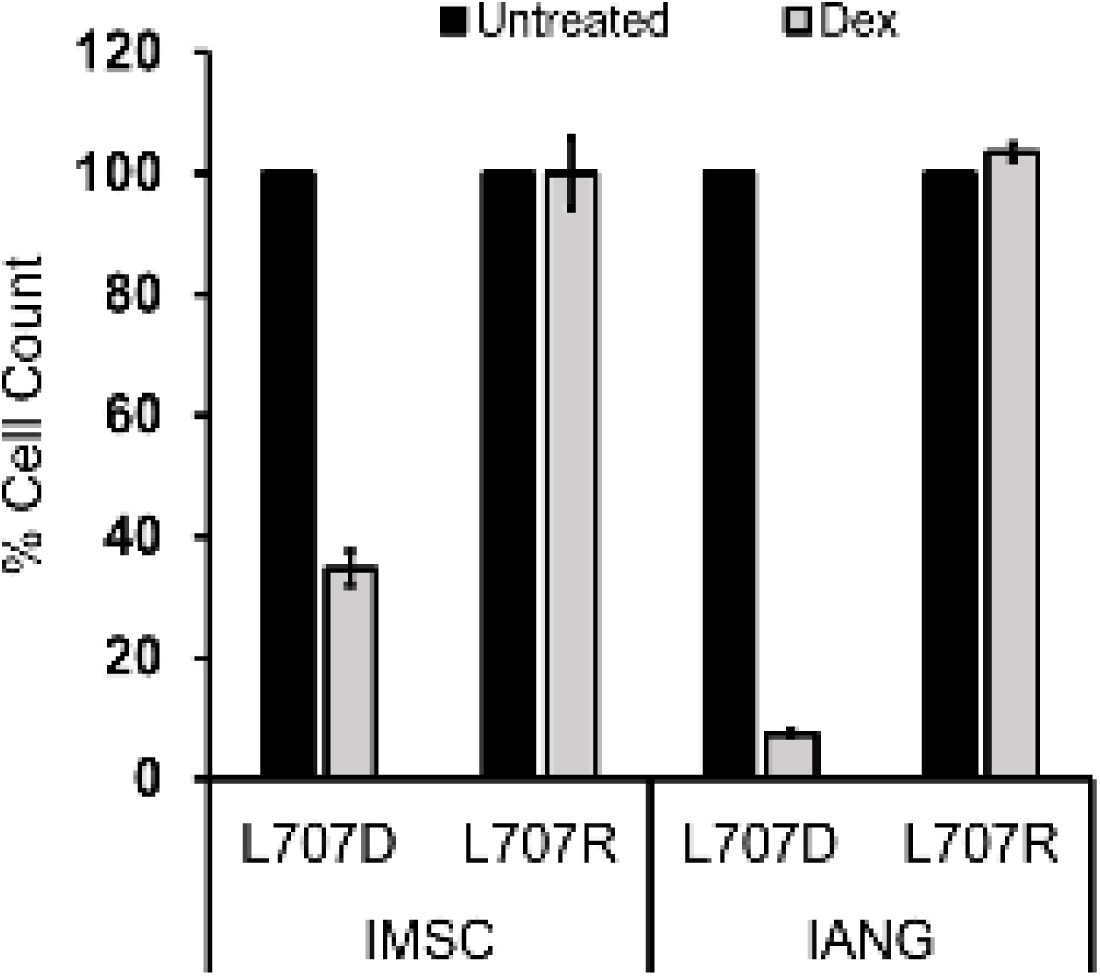
Cell counts following 10nM dexamethasone treatment on patient leukaemia cells at diagnosis [L707D] and relapse[L707R]. Counts taken at day 7

**S10.**
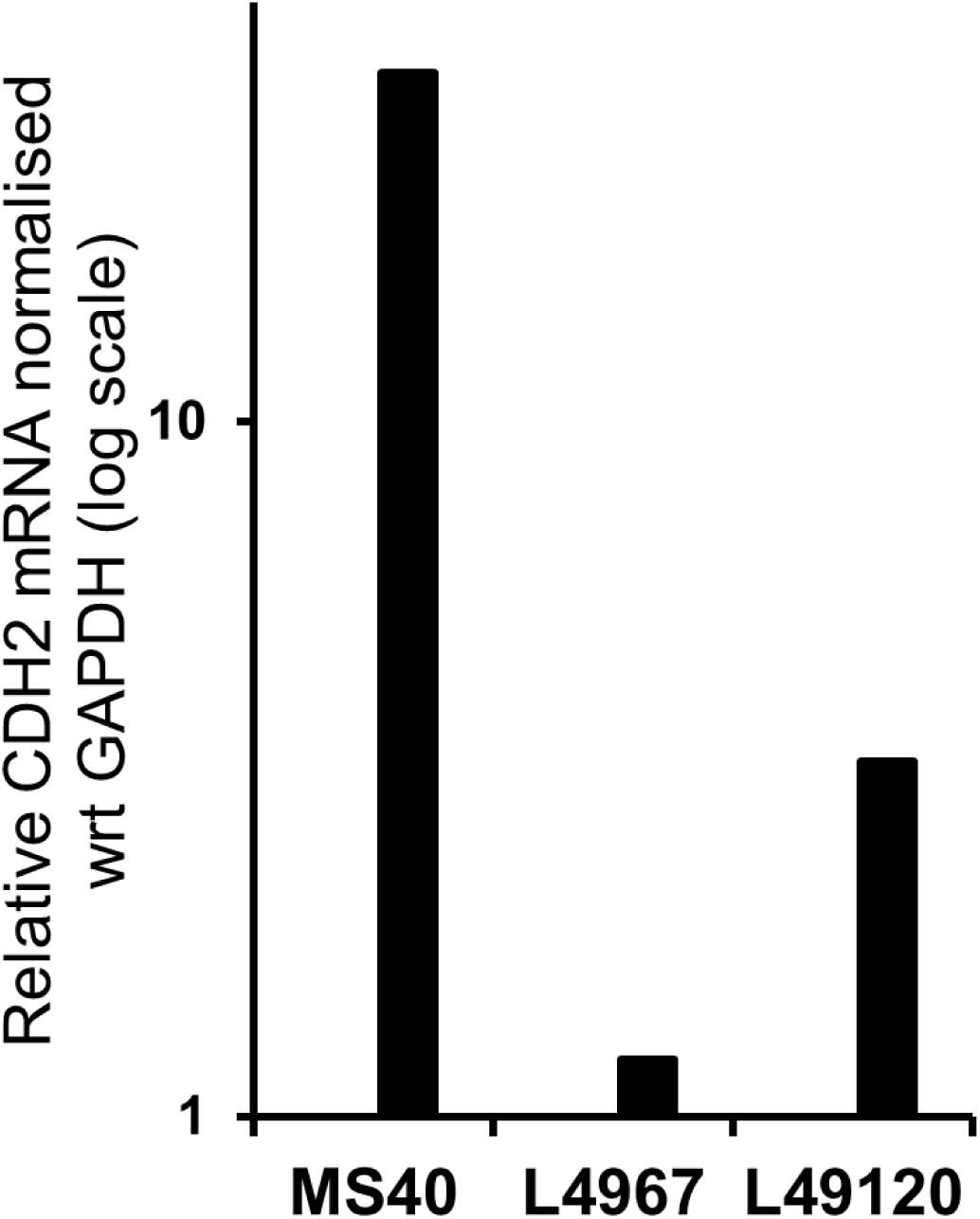
CDH2 mRNA levels in fast cycling iMSC-primed patient leukaemia cells (samples MS40, L4967, L49120) standardised against slow-cycling cells

**S11.**
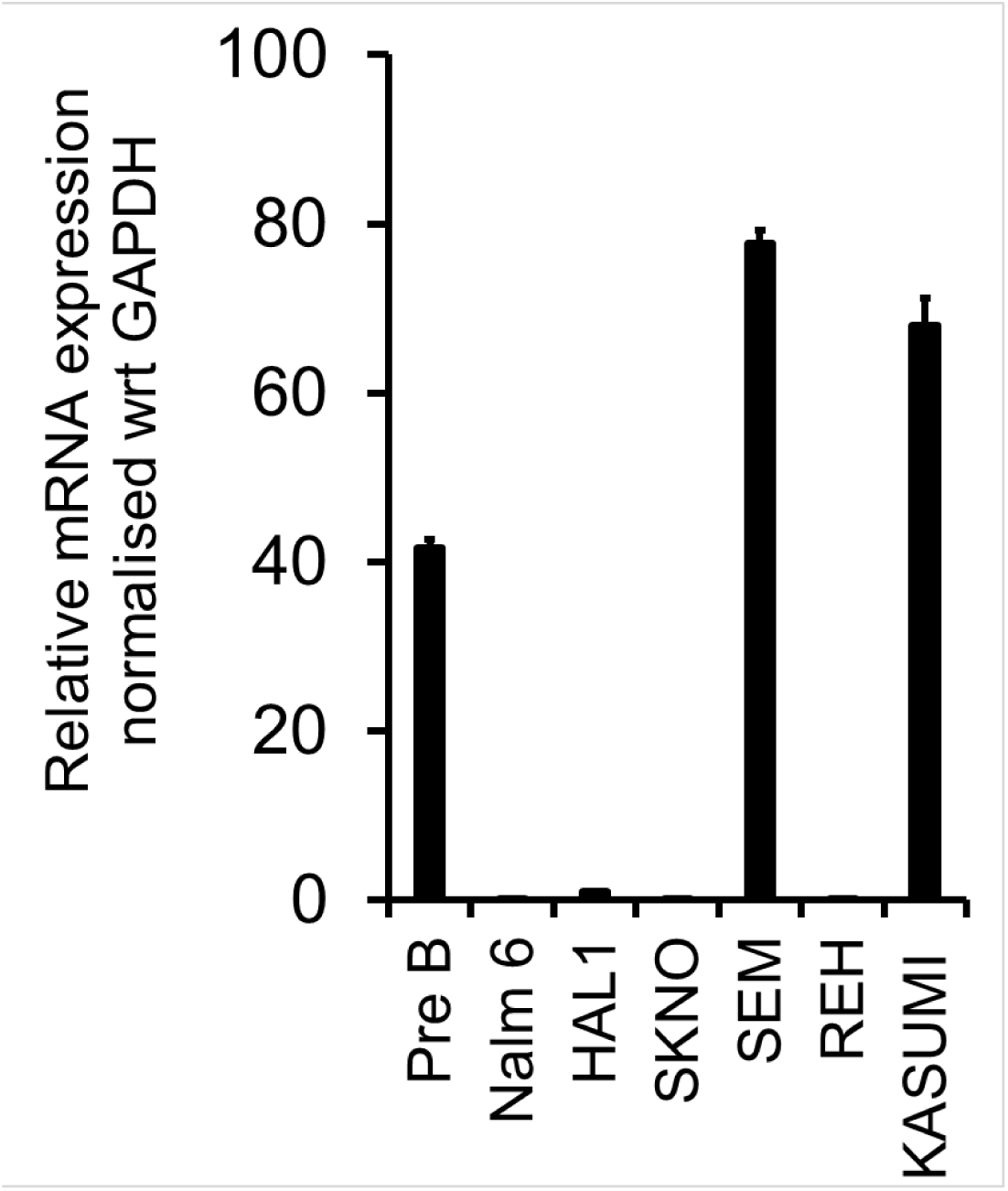
CDH2 mRNA levels in cell lines

**S12.**
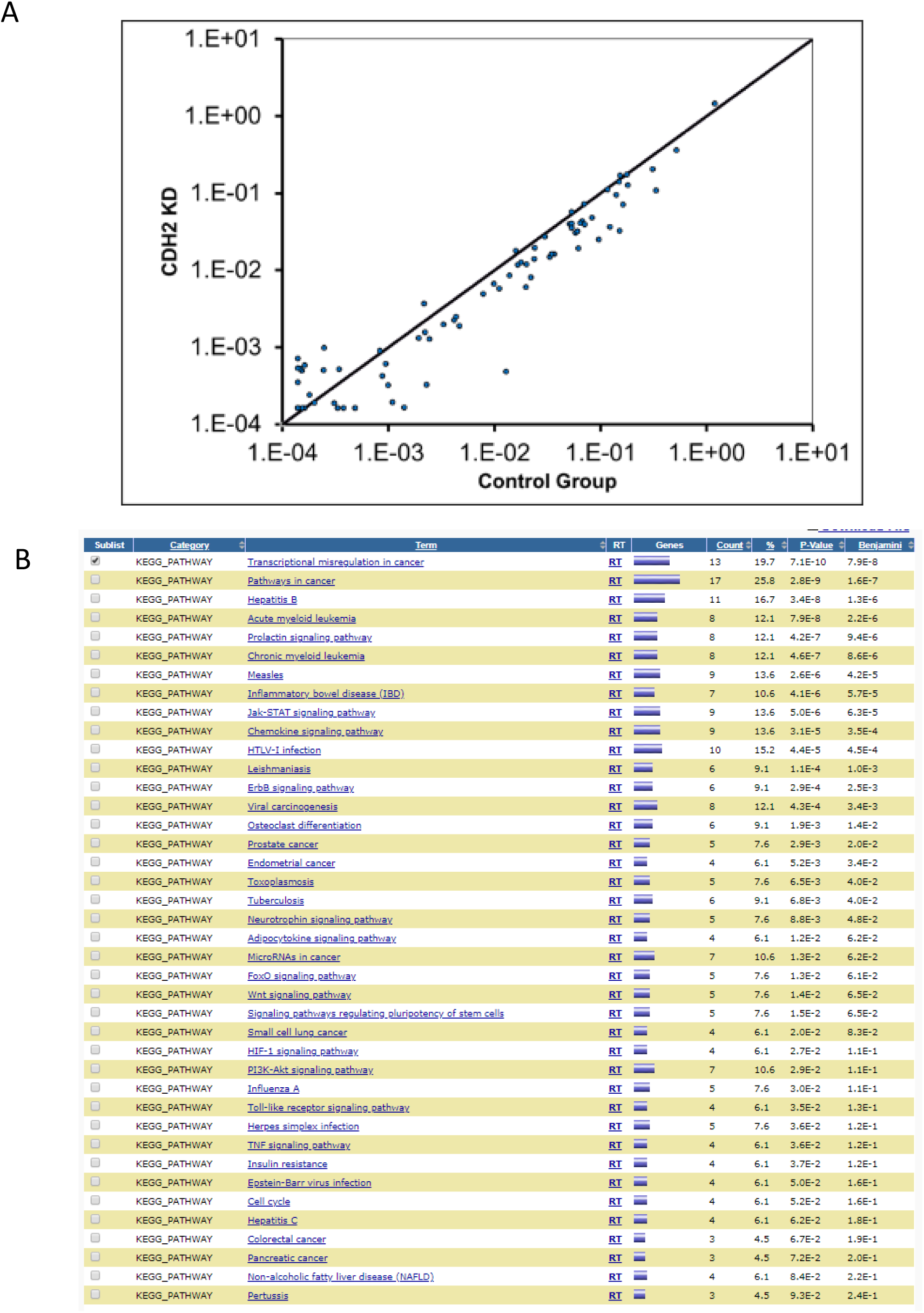
A. Scatter plot showing gene expression in control and CDH2 knockdown in SEM leukaemia cells. B. KEGG Pathways analysis in CDH2 knockdown in SEM leukaemia cells

**S13.**
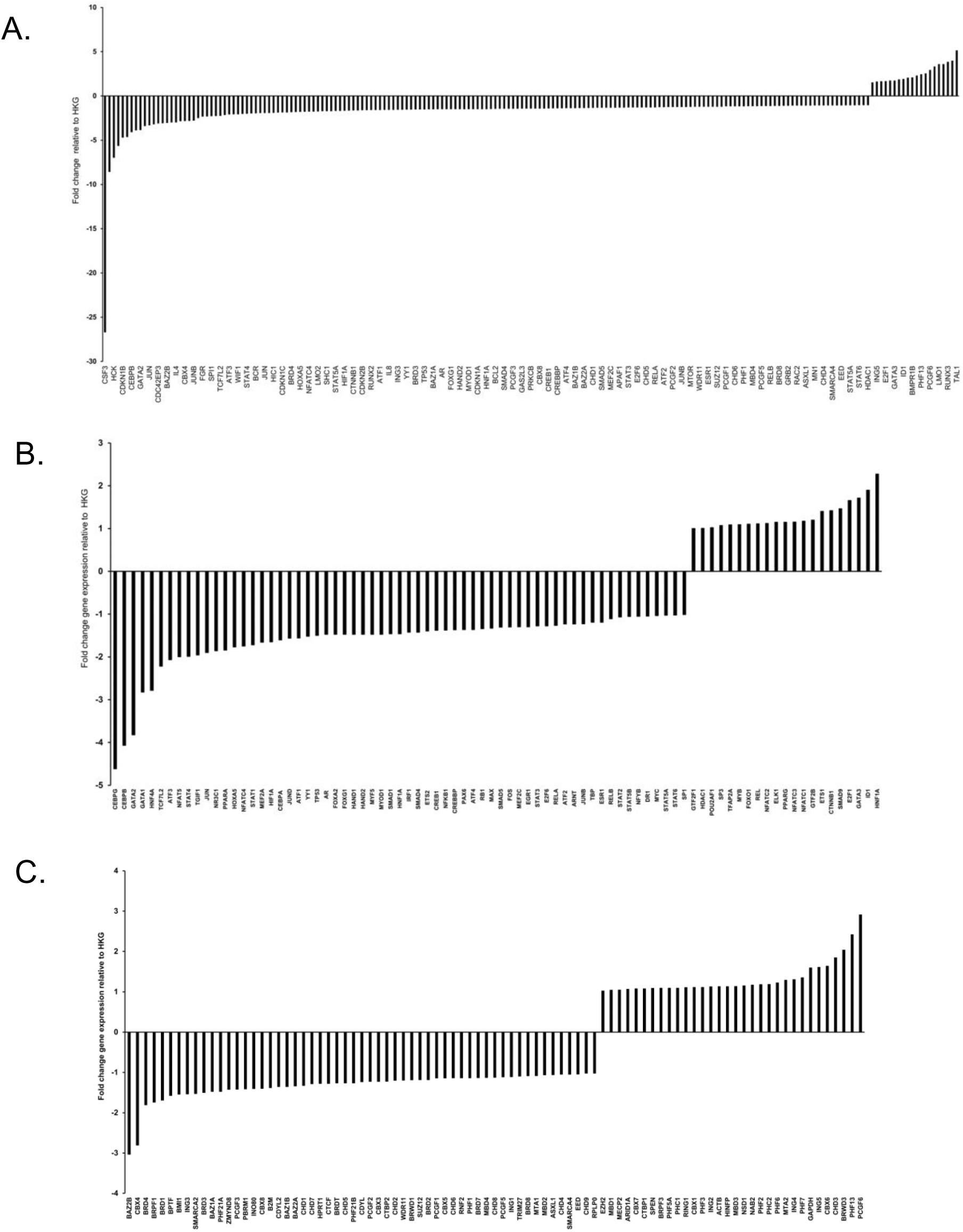
Fold change in gene expression following CDH2 shRNA knockdown in leukaemic cell line SEM. Genes profiled include A. those that play a role in human leukaemogenesis. B. Transcription factor and C. Chromatin remodelling factors

**S14.**
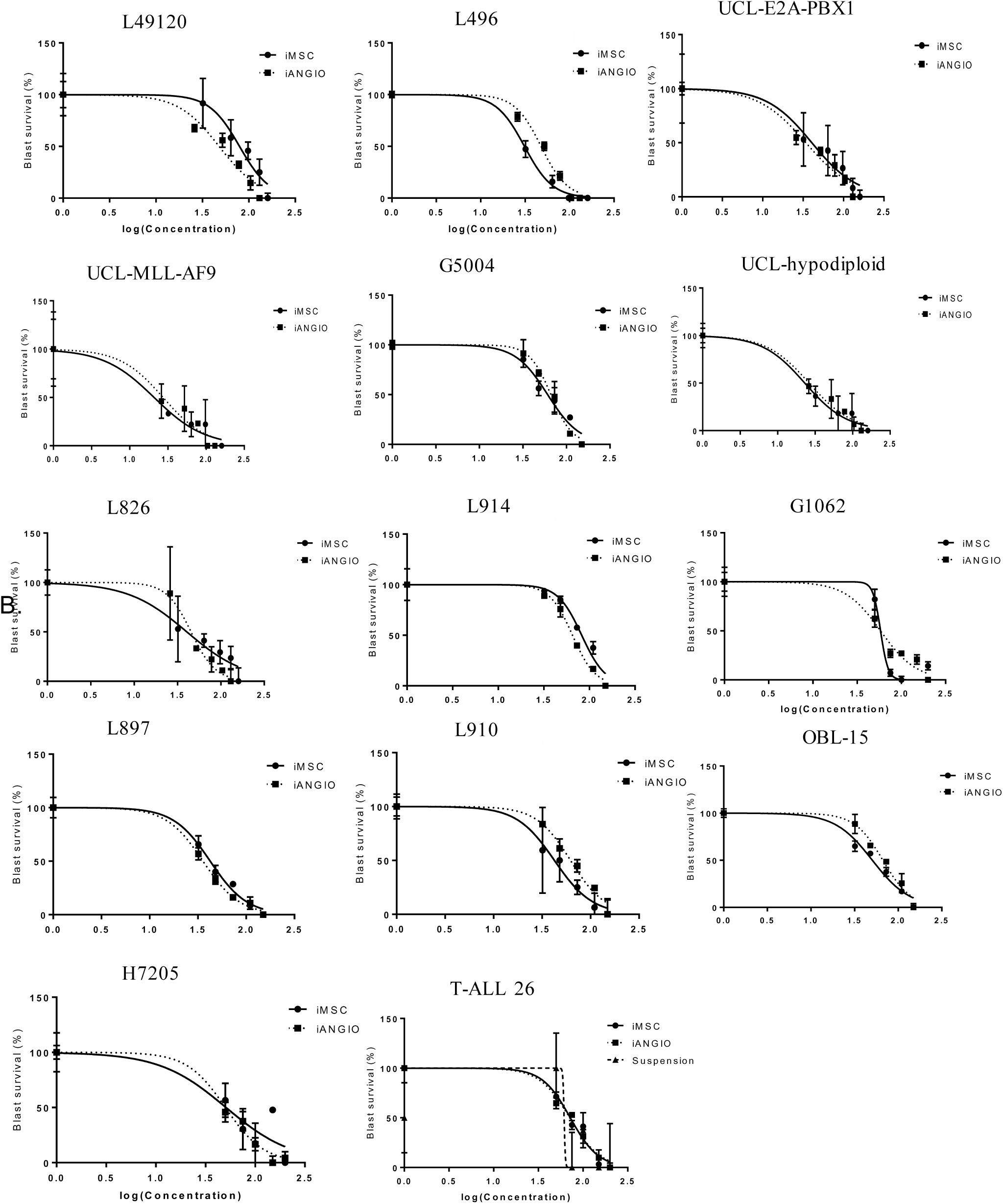
Drug dose response curves with ADH-1 (Exherin) on patient leukaemia samples

**S15.**
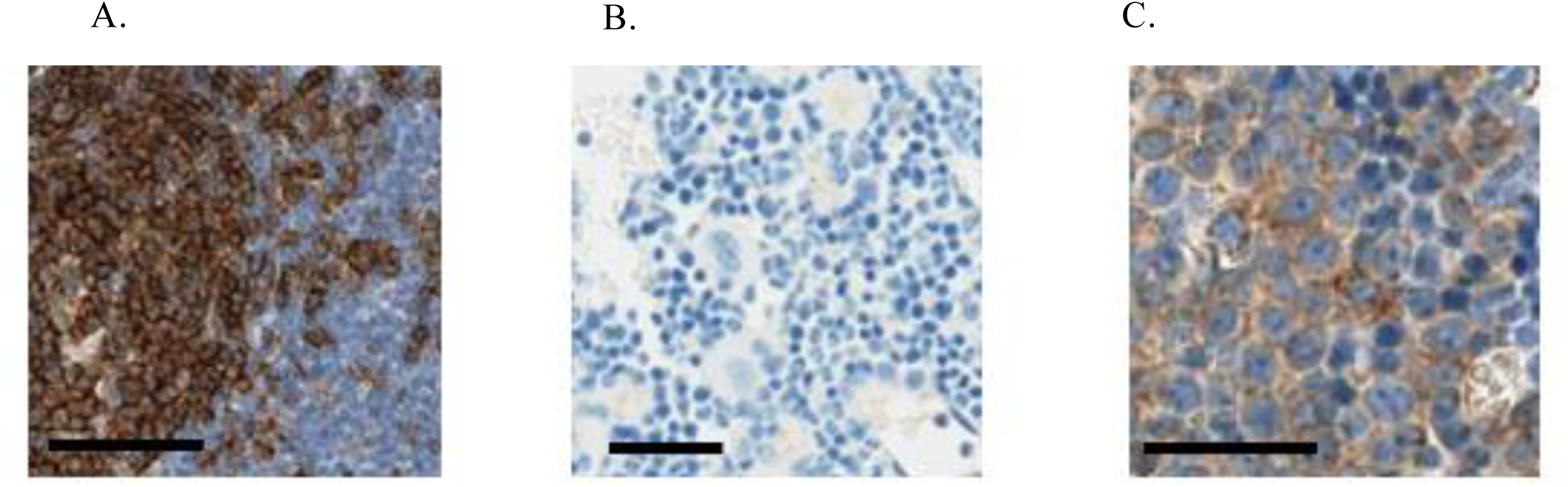
CD19 immunohistochemistry staining of sections from A. Human tonsil, positive control; B. Naïve, non-leukaemic mouse bone marrow, negative control, and C. Mouse bone marrow from a L707 PDX transplanted mouse at pre-treatment stage, with BLI total flux equivalent to efficacy study day6 pre-transplant. Scale bar = 50µm .

**S16.**
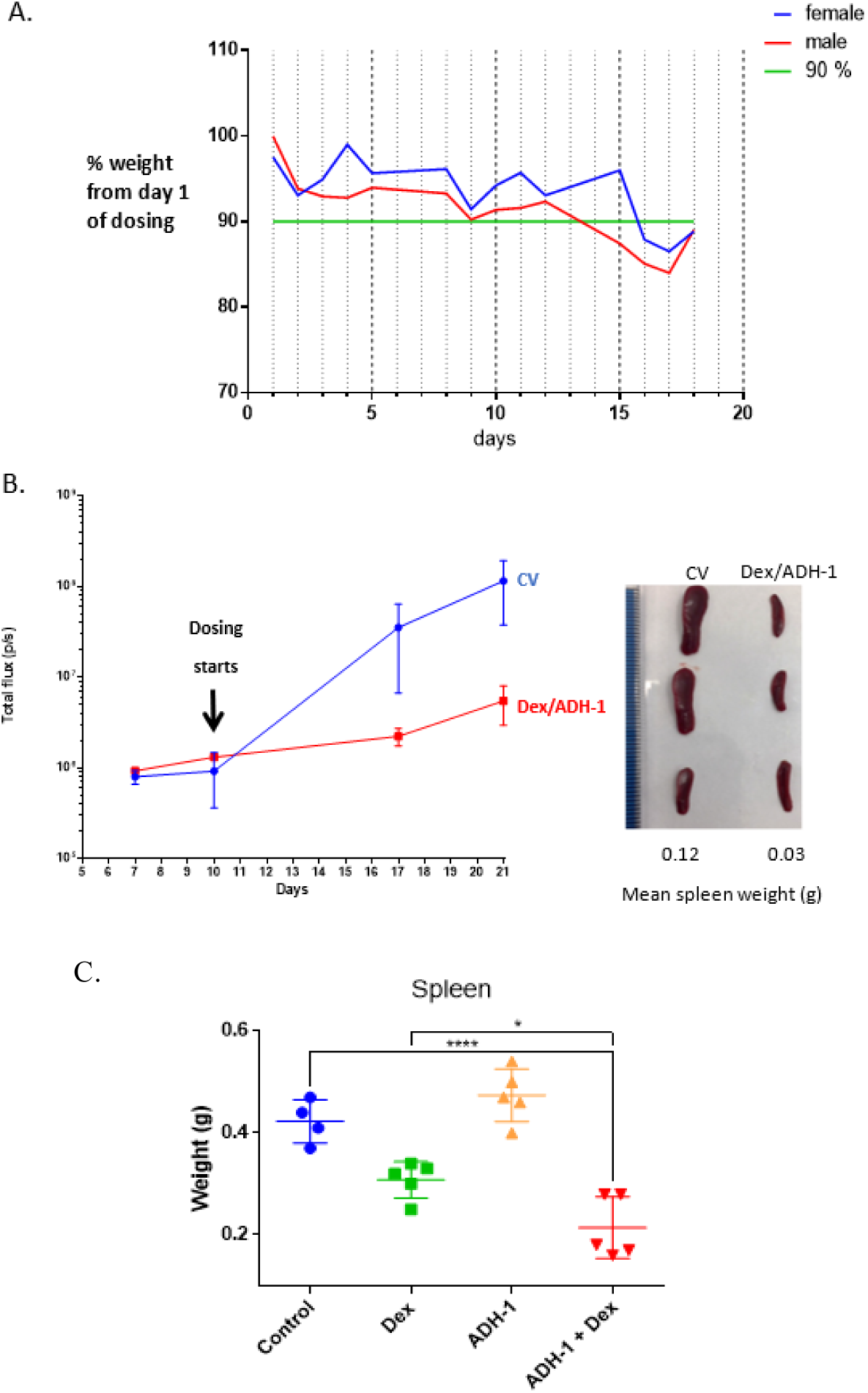
A. Weights of mice (2mice/sex) administered with 3mg/kg dexamethasone, 200mg/kg ADH-1 (Dex/ADH-1) via intraperitoneal injection, 1x daily, 5x weekly for 3 weeks. Mice appeared healthy with no clinical signs of ill health. Slight weight loss was observed consistent with single drug dexamethasone dosing. B. Dex/ADH -1 feasibility efficacy study. Mean bioluminescent imaging total flux (left) and spleens at 21 days (right) from L707D Luc+ ALL PDX mice, 3 mice/group treated with control vehicle (CV) or Dex/ADH-1 for 9 doses as for A. At 21 days the mean total flux is significantly different between CV and Dex/ADH-1 groups t-test, p=0.002.and spleen at 21 days are smaller. C. Spleen weights of mice treated as indicated. Lines indicate mean and SE, symbols are individual mice. 1 way ANOVA, * p<0.05, ****p<0.00005

## Supplementary tables

**Table S1.**
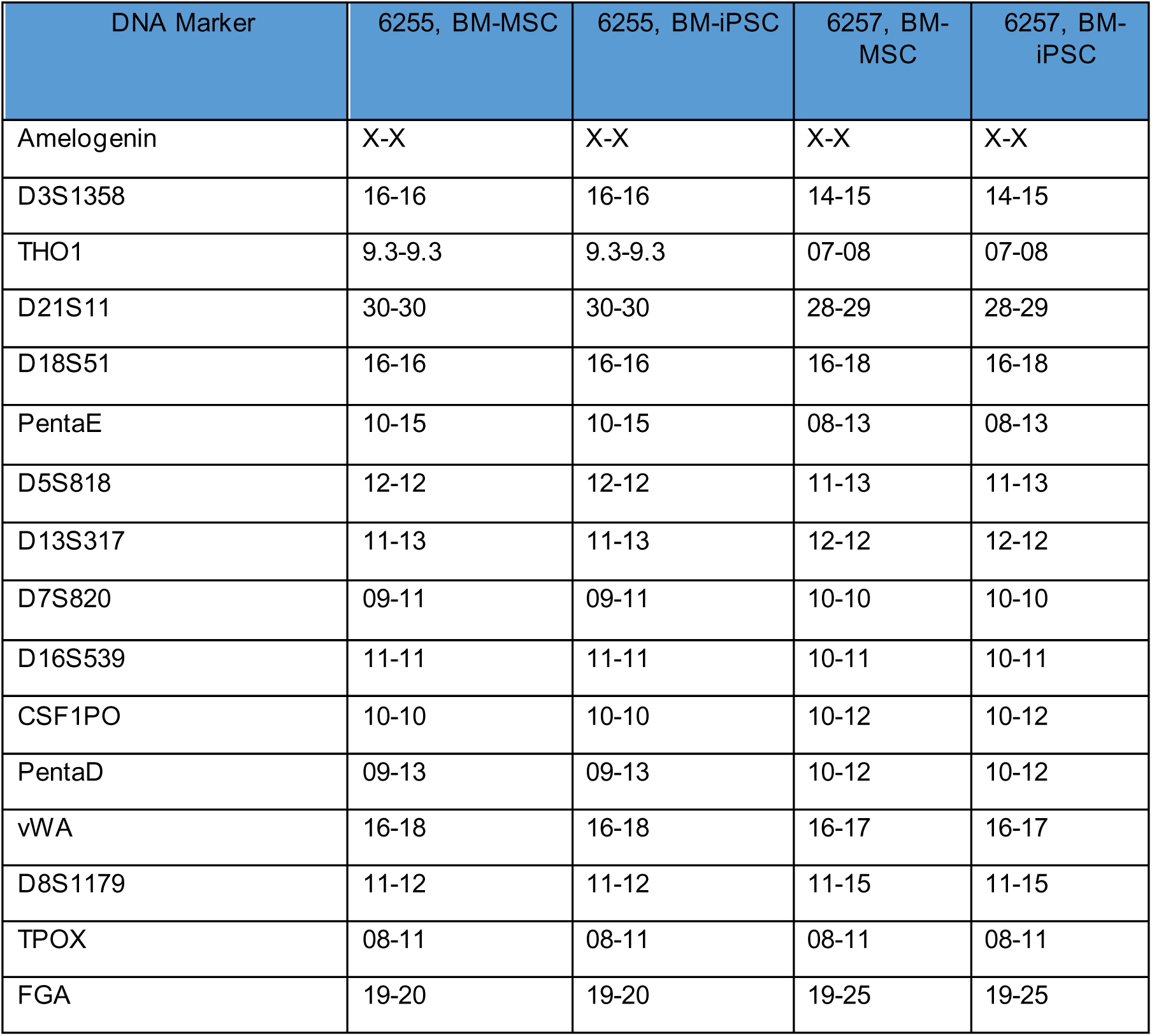
Microsatellite profiling confirms the BM-iPSC are an identical match to the parental primary bone marrow mesenchymal stroma cells for the 16 microsatellites tested including amelogenin, a sex marker

| DNA Marker | 6255, BM-MSC | 6255, BM-iPSC | 6257, BM-MSC | 6257, BM-iPSC |
| --- | --- | --- | --- | --- |
| Amelogenin | X-X | X-X | X-X | X-X |
| D3S1358 | 16-16 | 16-16 | 14-15 | 14-15 |
| THO1 | 9.3-9.3 | 9.3-9.3 | 07-08 | 07-08 |
| D21S11 | 30-30 | 30-30 | 28-29 | 28-29 |
| D18S51 | 16-16 | 16-16 | 16-18 | 16-18 |
| PentaE | 10-15 | 10-15 | 08-13 | 08-13 |
| D5S818 | 12-12 | 12-12 | 11-13 | 11-13 |
| D13S317 | 11-13 | 11-13 | 12-12 | 12-12 |
| D7S820 | 09-11 | 09-11 | 10-10 | 10-10 |
| D16S539 | 11-11 | 11-11 | 10-11 | 10-11 |
| CSF1PO | 10-10 | 10-10 | 10-12 | 10-12 |
| PentaD | 09-13 | 09-13 | 10-12 | 10-12 |
| vWA | 16-18 | 16-18 | 16-17 | 16-17 |
| D8S1179 | 11-12 | 11-12 | 11-15 | 11-15 |
| TPOX | 08-11 | 08-11 | 08-11 | 08-11 |
| FGA | 19-20 | 19-20 | 19-25 | 19-25 |

**Table S2.** Patient samples

| Sample Identification | Key molecular/clinical characteristics |
| --- | --- |
| L707D | E2A/HLF ; Sample at diagnosis |
| L707R | E2A/HLF; Sample at relapse |
| L49120 | BCR/ABL |
| L4951 | BCR/ABL |
| L4967 | BCR/ABL |
| UCL E2A/PBX1 | E2A/PBX1 |
| UCL MLL/AF9 | MLL/AF9 |
| UCL Hypodiploid | HYPODIPLOID |
| L826 | MLL/AF4 |
| LK124 | MLL/AF4 |
| G5004 | iAMP21 |
| G3131 | iAMP21 |
| G1062 | iAMP21 |
| H7205 | iAMP21 |
| G7578 | iAMP21 |
| L914 | HIGH HYPERDIPLOID |
| L897 | UNKNOWN |
| L910 | E2A/PBX1 |
| L590R | BCR/ABL |
| OBL-15 | HYPODIPLOID ADULT |
| T-ALL 26 | UNKNOWN |
| MS40 | BIPHENOTYPIC/MLL REARRANGEMENT |

**S3.**
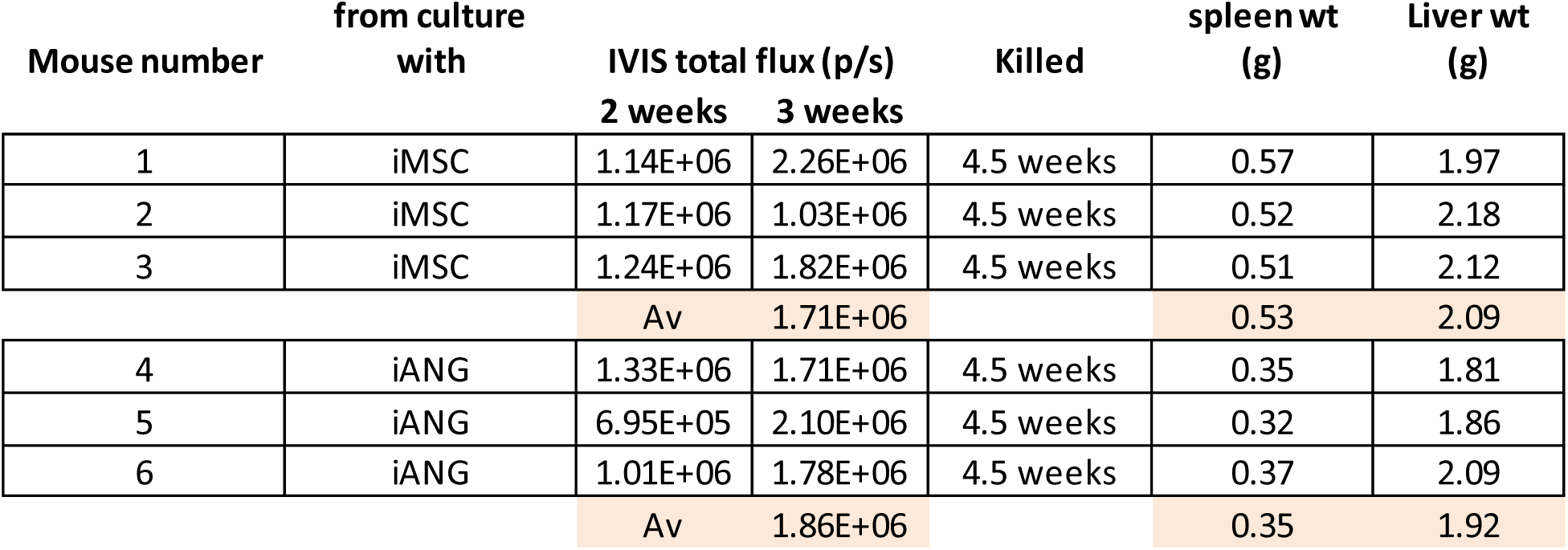
Bioluminescent imaging total flux and organ weights of NSG mice transplanted intrafemorally with 300,000 L707D PDX cells following their culture with i -niche. Total body engraftment (measured by total flux) did not differ significantly between the two niche grown PDX however the spleens of mice transplanted with iANG grown cells were significantly smaller (measured by weight) suggesting a reduction in system engraftment of cells to the BM. The liver in this PDX model contains very few engrafted cells so is a control for mouse size.

**Table S4.** Table of primary and secondary antibodies

| Marker | Primary antibody | Primary antibody<br>dilution factor | Secondary<br>antibody | Secondary<br>antibody dilution<br>factor |
| --- | --- | --- | --- | --- |
| Tra-1-60 | IgG3 mouse | 1:250 | Alexa Flour 594<br>goat anti-mouse | 1:500 |
| SSEA4 | IgM mouse | 1:250 | Alexa Fluor 488<br>anti-mouse | 1:500 |
| Sox2 | Alexa Fluor 647<br>anti-Sox2 | 1:250 |  |  |
| Oct4 | Alexa Fluor 488<br>anti-Oct4 | 1:250 |  |  |

## Acknowledgments

This study was funded by NC3Rs through a fellowship to DP, CCLG through a project grant to DP, JV and OH and a Wellcome Trust NUSCU award to DP. We would like to further thank CRUK for a programme grant C27943/A12788 to JV and OH and NECCR for funding core infrastructure at Wolfson Childhood Cancer Research Centre, Newcastle University. The IVIS Spectrum was funded by grant 087961 from the Wellcome Trust. We thank Universiti Sains Malaysia and Kementerian Pengajian Tinggi Malaysia for funding PhD studentship to AI. We also thank Dr. Malgorzata Firczuk, Medical University of Warsaw for the provision of leukaemia PDX samples.

## Funding

National Centre for the Replacement Refinement and Reduction of Animals in Research (NC3Rs) - NC/P002412/1 [DP]

Children’s Cancer and Leukaemia Group (CCLG) - CCLGA 2016 05 BH160568 [DP, JV, OH]

Wellcome Trust NUSCU award – OSR/0190/DPAL/NUSC [DP]

Cancer Research UK (CRUK) - C27943/A12788 [JV, OH]

## Autor contributions

Conceptualization: DP

Methodology: DP, HB, AF, JC

Investigation: DP, HB, SB, AHS, SN, AI, MB, RN, AW, MS, SS, RT, CK, AF, HM, LR, CS, PZ, PS

Formal analysis: SN

Visualization: DP, HB

Validation: DP, HB

Resources: DP, HB, JV, OH,

Funding acquisition: DP, JV, OH

Project administration: DP, HB

Supervision: DP, HB, JV, OH

Writing – original draft: DP, HB

Writing – review & editing: DP, JV, OH, AM, CJH, JMA, CH

## Data and materials availability

All data are available in the main text or the supplementary materials

